# miRNAs and neural alternative polyadenylation specify the virgin behavioral state

**DOI:** 10.1101/2020.04.05.026831

**Authors:** Daniel L. Garaulet, Binglong Zhang, Lu Wei, Elena Li, Eric C. Lai

## Abstract

How are diverse regulatory strategies integrated to impose appropriately patterned gene expression that underlie *in vivo* phenotypes? Here, we reveal how coordinated miRNA regulation and neural-specific alternative polyadenylation (APA) of a single locus controls complex behaviors. Our entry was the unexpected observation that deletion of Bithorax-Complex (BX-C) miRNAs converts virgin female flies into a subjective post-mated behavioral state, normally induced by seminal proteins following copulation. Strikingly, this behavioral switch is directly attributable to misregulation of *homothorax* (*hth*). We localize specific CNS abdominal neurons where de-repressed Hth compromises virgin behavior in BX-C miRNA mutants. Moreover, we use genome engineering to demonstrate that precise mutation of *hth* 3’UTR sites for BX-C miRNAs, or deletion of its neural 3’ UTR extension containing most of these sites, both induce post-mated behaviors in virgins. Thus, facilitation of miRNA-mediated repression by neural APA is required for virgin females to execute behaviors appropriate to their internal state.

## Introduction

Whether to sleep when tired, drink when thirsty, or eat when hungry, reproductive and survival success depends on the capacity for dynamic coordination of internal states with appropriate behaviors. A classic example involves the profound behavioral transformations that females of many species undergo following mating and pregnancy. Mature virgins typically exhibit receptivity to male courtship and eventually allow copulation. However, after mating, they tend to remain refractory to further copulation attempts and adapt their habits to produce, raise and protect their progeny (Clyne and Miesenbock, 2009). This behavioral transition is found across diverse phyla, from nest building or increased aggression in mice (Broida and Svare, 1982; Ogawa and Makino, 1984), to a collective behavioral remodeling in insects (Avila et al., 2011). Overall, as a result of mating or pregnancy, diverse behaviors of an inseminated female adapt to match the novel needs of the reproductive state.

Alterations of female behaviors induced by mating have been extensively studied in female fruitflies, where multiple reproductive conducts such as egg-laying and male acceptance (Connolly and Cook, 1973; Manning, 1962, 1967), but also appetite (Carvalho et al., 2006), lifespan (Fowler and Partridge, 1989), food and salt preference (Ribeiro and Dickson, 2010; Walker et al., 2015), immune system function (Peng et al., 2005b), and sleep (Isaac et al., 2010) are modulated by sexual interaction. Collectively, these behaviors comprise the post-mating switch, also referred to as the post-mating response (PMR) (Avila et al., 2011). The capacity of a male fly to manipulate female behavior resides within seminal fluids synthesized by the male accessory gland (Gillott, 2003). While many seminal fluid proteins are known (Avila et al., 2011), the 36 amino acid Sex peptide (SP) is sufficient to drive most female post-mating responses (Aigaki et al., 1991; Chen et al., 1988). SP associates with sperm and is transferred to the female genital tract during copulation (Peng et al., 2005a). It is then stored in specific female reproductive organs and released gradually, thereby conditioning female behavior for 8-10 days after copulation (Manning, 1962, 1967; Peng et al., 2005a). Considering the average fruitfly lifespan is 30-40 days, it is remarkable that an exogenous factor, transferred in a single act, can drive female behavior for ¼ of her life.

SP exerts its function through a G-coupled transmembrane receptor, SPR (Yapici et al., 2008). Although SPR is expressed throughout the abdominal ganglion of the ventral nerve cord (VNC, equivalent of the vertebrate spinal cord) and in some regions of the brain (Yapici et al., 2008), its expression in a restricted set of 6-8 *pickpocket*+ sensory neurons (SP sensory neurons, SPSN) adjacent to the uterus appears necessary and sufficient to drive most effects of the switch (Feng et al., 2014; Hasemeyer et al., 2009; Yang et al., 2009). While some of these cells send their processes directly to the subesophageal ganglion in the female brain, the majority of SPSN terminations target the abdominal ganglion of the VNC (Feng et al., 2014; Hasemeyer et al., 2009; Yang et al., 2009), where they contact their synaptic partners. Some of these are abdominal interneurons that express myoinhibitory peptide (Mip) (Jang et al., 2017), which input into a restricted population of ascending neurons (SAG) that project to the posterior region of the brain (Feng et al., 2014; Soller et al., 2006). SPSN-Mip-SAG define a minimal ascending axis through which information flows from the SP stored in the genital organs to the female brain. However, beyond this simple, linear afferent circuit, larger populations of less characterized neurons are also involved either in the interpretation of mating information or the execution of mated responses (Haussmann et al., 2013). Most of these neurons express *fruitless*, *doublesex* and/or *pickpocket* genes, and configure an intricate neuronal network underlying female sexual behavior (Bussell et al., 2014; Feng et al., 2014; Haussmann et al., 2013; Monastirioti, 2003; Rezaval et al., 2012).

Nonetheless, despite intense investigations of the behavioral, neuronal, and molecular nature of the post-mated switch, the genetic determinants of the virgin state remain largely unknown. Here, we report that developmental repression of an individual transcription factor by microRNAs (miRNAs) is critical to establish virgin behavior. This regulatory axis was revealed by deletion of the bidirectionally transcribed Bithorax Complex (BX-C) miRNA locus, which constitutively induce multiple post-mated behaviors in virgins. The BX-C hairpin locus expresses distinct miRNAs (miR-iab-4/miR-iab-8) in adjacent domains of the ventral nerve cord, which contains specific neurons that mediate the post-mating switch. Deletion of the BX-C miRNAs strongly derepresses *homothorax* (*hth*), which is directly targeted by both BX-C miRNAs to suppress post-mated behavior in virgins. Importantly, targeted mutation of the array of BX-C miRNA binding sites in the *hth* 3’ UTR similarly abrogates virgin behavior. Finally, we integrate this with the regime of neural alternative polyadenylation, since specific deletion of the *hth* neural 3’ UTR extension that bears most of these miRNA binding sites also blocks the virgin state.

Overall, we utilize miRNAs as an entry point to reveal critical post-transcriptional biology that interfaces with alternative polyadenylation control. The failure of these regulatory events has profound consequences on the ability of female flies to integrate sexual internal state with appropriate behaviors.

## Results

### BX-C miRNAs suppress the post-mating switch in virgins

Egg-laying is a defining characteristic of mated behavior in female flies. Although old virgins are able to lay some unfertilized eggs, oviposition is largely suppressed in young (0-3 day old) virgins (**Figure 1A** and **Supplementary Figure 1B, C**). Instead, mating and SP reception induces females to lay eggs (Manning, 1962, 1967). Surprisingly, young virgin females deleted for the bidirectionally transcribed BX-C miRNA locus encoding miR-iab-4 and miR-iab-8 (*mir[*Δ]) exhibit inappropriate, robust, egg-laying activity (**Figure 1A**). Since many behavioral phenotypes are influenced by unknown genetic background factors, we used CRISPR/Cas9 to generate a new BX-C miRNA deletion (*mir[C11]*, **Supplementary Figure 2**), and extensively backcrossed both alleles into a common wildtype reference strain (*Canton-S*). We consistently observed activation of egg-laying in all BX-C miRNA deletion allele combinations in comparison to multiple wildtype strains (**Figure 1A** and **Supplementary Figure 1B,C**).

**Figure 1.**
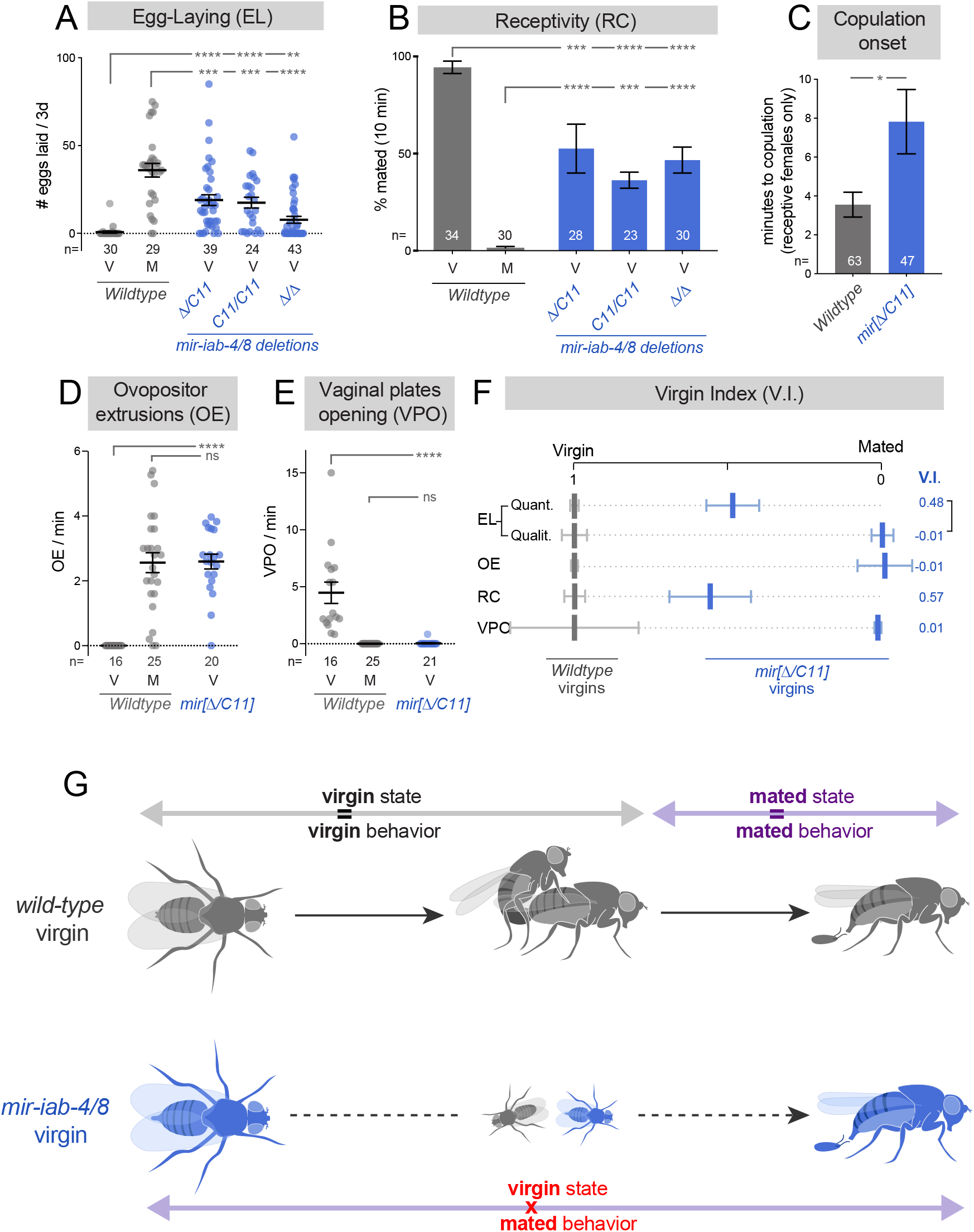
BX-C miRNAs prevent post-mating behaviors in virgins. (A-E) Deletion of mir-iab-4/8 suppresses multiple virgin behaviors (B,C,E) and induces mated behaviors (A,D) in virgin females. (A) Quantification of egg-laying by individual females that are either virgin (V) or mated (M). (B) Analysis of sexual receptivity, assessed by the fraction of individual females that are copulating at 10 minutes across multiple genotypes. (C) Timing of copulation onset, analyzed in the subset of females that are receptive to mating. (D) Quantification of ovipositor extrusions by individual females. Virgins are 1 day old. (E) Quantification of vaginal plates openings by individual females. (F) A metric summarizing all of the behaviors studied (virgin index), showing that deletion of *mir-iab-4/8* shifts virgin behavior to mated values. Egg-laying is presented in both quantitative (Quant.) and qualitative (Qualit., yes/no) measures. (G) Schematic representation of internal state and behavior in wildtype (gray) and BX-C miRNA mutant females (blue). While behavior is constantly coupled to mating state in wildtype females, deletion of *mir-iab-4/8* in virgins suppresses virgin behaviors and triggers post-mating responses before copulation. Statistical significance was evaluated using Fisher’s exact test for receptivity (B), and Mann-Whitney non parametric test for all other behaviors. ns=not significant, * p<0.05, ** p<0.01, *** p<0.001, **** p<0.0001. Error bars= SEM. Wildtype flies are *Canton-S* strain.

Of note, mated *mir[*Δ] females were previously documented to have defective egg-laying, which was previously speculated to be due to egg blockage (Bender, 2008; Gummalla et al., 2012) and/or reduced oviduct innervation (Garaulet et al., 2014). However, the robust oviposition by young virgins indicates functional neuromuscular control at the genital tract, and instead suggests a subjective shift to the mated state. Therefore, we sought evidence for alterations in other switch-influenced behaviors in miRNA mutant female virgins.

We first assayed female receptivity, which is maximal in mature virgins and allows the male to copulate shortly after courtship initiation. Receptivity is strongly suppressed following mating, as females remain refractory to male copulatory attempts for days (Connolly and Cook, 1973) (**Figure 1B** and **Supplementary Figure 1A**). *mir-iab-4/8* null virgins show compromised receptivity across all mutant combinations tested, evaluating both mating success and time elapsed until copulation in the subset of females that eventually mate (**Figure 1B,C**, **Supplementary Figure 1A, Supplementary Figure 3** and **Supplementary Movies 1 and 2**).

Male acceptance or rejection is achieved by distinct female genital responses, which change according to the female mating state. Mature virgins typically open their vaginal plates sporadically, while mated females extrude their ovipositor to reject male copulatory attempts (Aranha and Vasconcelos, 2018; Bussell et al., 2014; Connolly and Cook, 1973). For both of these behaviors, *mir[*Δ*/C11]* virgins were indistinguishable from mated wildtype flies (**Figure 1D, E** and **Supplementary Figure 1D, E**). Given the reliability of these different phenotypic readouts, we focused subsequent analyses of the miRNA null state using trans-heterozygous deletions (*mir[*Δ*/C11]*).

Taken together, the overall performance of BX-C miRNA knockout virgins is shifted to a canonical post-mated response (**Figure 1F,G**), indicating that miR-iab-4/8 are crucial for the subjective interpretation of internal state by virgin females. Importantly, this is not only revealed by the failure to perform characteristic virgin behaviors, but also by the selective activation of behavioral programs that are solely executed by mated flies (**Figure 1F,G**). Moreover, while virgin behaviors are commonly presumed to be a default state that must be converted to the mated state, the loss of BX-C-miRNAs reveal that virgin behavior is genetically specified.

### BX-C miRNAs are expressed in VNC subpopulations that control the post-mating switch

Behavior on either side of the switch is determined by an ascending neural circuit that underlies SP sensing. Although the precise architecture of the circuit has not been fully mapped, most of its neuronal substrates reside in the abdominal ganglion (AbG) of the ventral nerve cord (VNC), and can be identified by expression of *fruitless* (*fru*), *pickpocket* (*ppk*) or *doublesex* (*dsx*) genes (Feng et al., 2014; Hasemeyer et al., 2009; Jang et al., 2017; Rezaval et al., 2012; Yang et al., 2009). While these are broadly expressed markers, some subsets of more restricted neuronal populations labeled by select VT-Gal4 lines (hereafter VT-switch neurons) have also shown to be necessary and sufficient to fully induce post-mating behaviors (Feng et al., 2014). We studied how BX-C miRNAs are deployed with respect to these neural subpopulations. These miRNAs are expressed in adjacent domains within the developing abdominal ganglion of the larval VNC, with miR-iab-4 deployed anteriorly to miR-iab-8 (Bender, 2008; Garaulet et al., 2014; Garaulet and Lai, 2015; Tyler et al., 2008). In the VNC of adult females, we used tub-GFP-miR-iab-4 and miR-iab-8 sensors to register the domains of miRNA activity with respect to Abd-A protein (**Figure 2A, B**).

**Figure 2.**
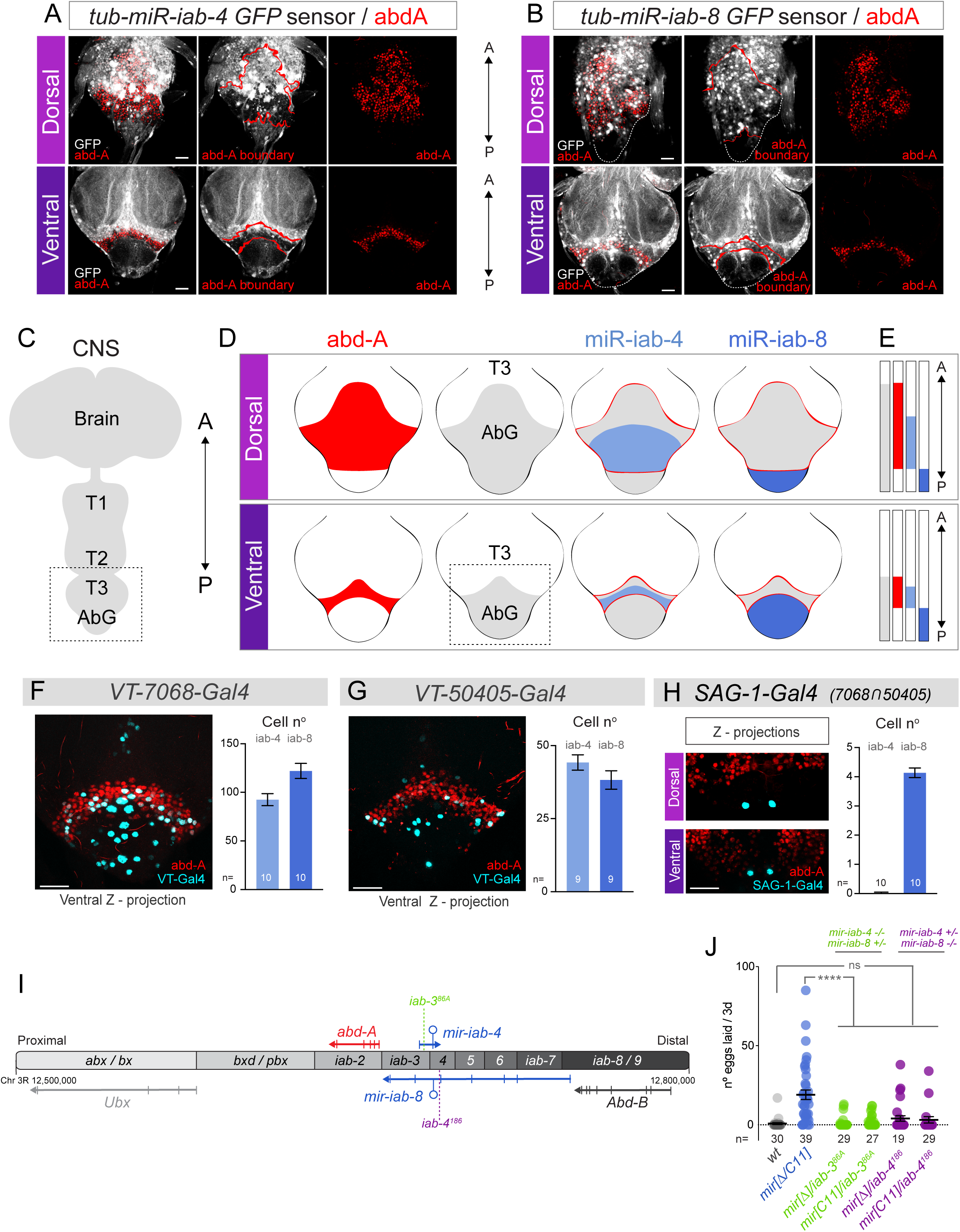
BX-C miRNA functional domains in the VNC. Shown are immunostainings of the *Drosophila* adult VNC, focused on the abdominal ganglion (AbG, box in C). (A,B) *tub-GFP* activity sensors to reveal domains of active miR-iab-4 (A) and miR-iab-8 (B), as inferred from reduced GFP accumulation. Co-staining with Abdominal-A (red) reveals collinear, abdominal-exclusive activity of both miRNAs along the A/P axis. (C) Schematic of the *Drosophila* CNS. Boxed region corresponds to T3-AbG shown in A, B and D. (D, E) Abdominal patterning of the VNC. Although the size of each domain differs in the dorsoventral axis, the relative A/P positioning of each domain is maintained. Boxed region in (D) corresponds to the ventral side of AbG shown in F-H. (F-H) Expression pattern and quantification of VT-switch lines. Co-expression with Abd-A reveals an enrichment of Gal4+ cells in the AbG, within the domains of expression of *mir-iab-4* and *mir-iab-8*. (I) Diagram of the Bithorax-Complex. *abx*=*anterobithorax*, *bx*=*bithorax*, *bxd*= *bithoraxoid*, *pbx*=*postbithorax*, *iab*=*infra-abdominal*, *Ubx*=*Ultrabithorax*, *abd-A*=*abdominal-A*, *Abd-B*=*Abdominal-B*. Note that the *mir-iab-4* and *mir-iab-8* hairpins are encoded by the same genomic DNA, but are transcribed from the top and bottom strands of the BX-C, respectively. (J) Egg-laying performance of individual *mir-iab-4* (*iab3^86A^/ mir[*Δ], green) or *mir-iab-8* mutants (*iab-4^186^/ mir[*Δ], purple). A=anterior, P=posterior, CNS=central nervous system, AbG=abdominal ganglion, T= thoracic segments. Error bars= SEM. Scale bar= 25μm. *mir-iab-4/8−/−* mutants=*mir[*Δ*/C11]* transheterozygotes.

In these analyses, it is relevant to consider the three-dimensional structure of the VNC, as the domains of these individual markers are substantially shifted between the dorsal and ventral surfaces. Still, the relative positions of Abd-A and the miRNAs are preserved between the dorsal and ventral regions of the adult VNC. Abd-A is present anterior to, and within the domain of active miR-iab-4, while active miR-iab-8 is exclusively posterior to Abd-A (**Figure 2A-E**). These relative relationships also apply in the embryo and larva (Bender, 2008; Garaulet et al., 2014; Gummalla et al., 2012; Tyler et al., 2008). Thus, we can use Abd-A (1) as a proxy for the transition of the two miRNA domains, and (2) to demarcate the abdominal ganglion.

We used co-labeling to define the domains of the VT-switch lines with respect to the BX-C miRNAs (**Figure 2F-H**). The most restricted driver, the split-Gal4 combination *SAG-1*, which derives from the regulatory intersection of the *VT-7068* and *VT-50405* lines, labels only four neurons that reside strictly within the miR-iab-8 domain (**Figure 2H**). However, the remainder of VT-switch lines are active in both miR-iab-4 and miR-iab-8 functional domains. We quantified the number of neurons labeled by each VT-switch Gal4 line in the respective miRNA domains in **Figure 2F-H** and **Supplementary Figure 4**. These data suggest that both miRNA loci might be involved in maintaining virgin state.

To test this notion, we separated the function of the bidirectional miRNA locus by placing a deletion allele in trans to chromosomal rearrangements that disrupt the primary transcripts for either top or bottom strand loci (**Figure 2I**). This approach was used to provide evidence that certain BX-C miRNA deletion phenotypes rely primarily on *mir-iab-4* (larval self-righting) (Picao-Osorio et al., 2015) or *mir-iab-8* (sterility) (Bender, 2008), respectively. We focused on egg-laying to readout the post-mating switch in representative breakpoint hemizygote backgrounds. While this behavior was sporadically activated in null conditions of either miRNA, neither was significantly different from control (**Figure 2J**). Thus, it appears that both miR-iab-4 and miR-iab-8, and by extension, neurons resident in both of their expression domains, contribute to the virgin behavioral defect in miRNA deletion females.

### Loss of BX-C miRNAs affects the activity of post-mating switch neurons

Several studies demonstrate that virgin behavior is determined by a constitutive activation of abdominal networks of neurons identified by the *fru*, *ppk*, *dsx* or VT-switch markers. In particular, hyperactivation of these neurons renders virgin behaviors in mated flies (Feng et al., 2014; Hasemeyer et al., 2009; Jang et al., 2017; Rezaval et al., 2012; Yang et al., 2009). As loss of miR-iab-4/8 induces post mating responses, we hypothesized that enforced neural activity might revert these conducts back to virgin state in *mir-iab-4/8* knockouts.

Following previous studies (Feng et al., 2014), we used the bacterial sodium channel transgene *UAS-NachBac* to enhance the activity of VT-switch neurons. We recapitulate prior behavior observations with the switch-Gal4 lines (**Figure 3A**) and show that our miRNA mutants are not rescued by inclusion of the *UAS-NaChBac* alone, ruling out leaky transgene effects (**Figure 3B**). Strikingly, activation of either *VT-7068* or *VT-50405* neurons substantially suppressed oviposition by miRNA deletion virgins, with *VT-7068* providing near complete rescue (**Figure 3C**). Activation of *VT-454* neurons also suppressed virgin oviposition in BX-C miRNA deletions (**Supplementary Figure 4B**).

**Figure 3.**
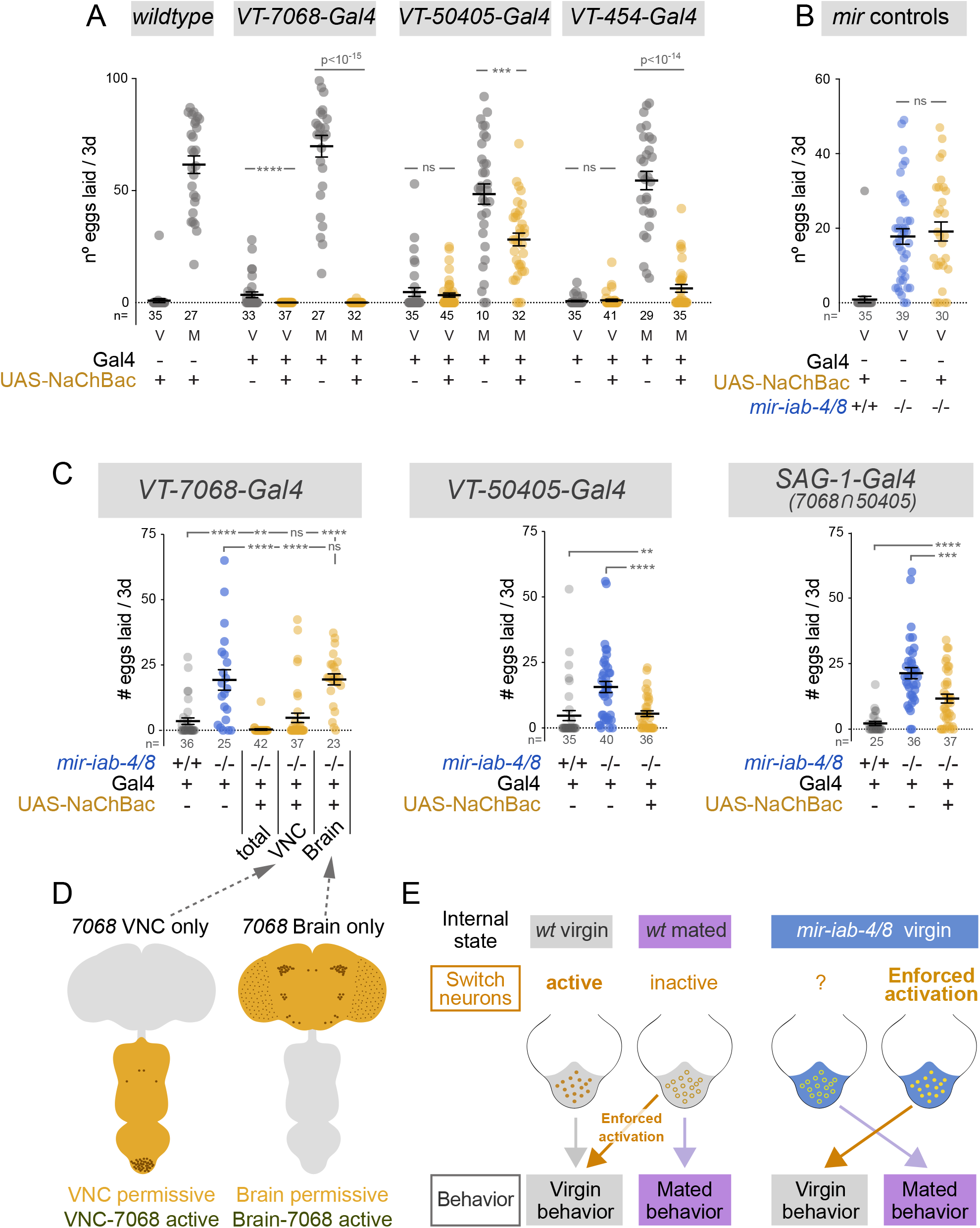
Enforced activation of abdominal neurons restores virgin behavior in *mir-iab4/8* mutants. (A) Hyperactivation of VT-switch neurons in wildtype mated females reverts egg-laying to virgin values. (B) *UAS-NaChBac* control in mir-iab-4/8 null background. (C) Activation of multiple VT-switch neurons in *mir-iab-4/8* mutant virgins suppressed their egg-laying phenotype. Activation of VNC but not Brain VT-7068 neurons alone accounts for the behavioral rescue. (D) Illustration of the VNC VS brain specific NaChBac activation of *VT-7068* neurons achieved in the following genotypes: *OTD-flp, tub>stop>Gal80 /+; 7068-Gal4, mir-C11/ UAS-NaChBac,* Δ*mir* and *OTD-flp, >tub-Gal80> /+; 7068-Gal4, mir-C11/ UAS-NaChBac,* Δ*mir*. (E) Schematic representation of the effect of neuronal activity and activity manipulations in wildtype and *mir-iab-4/8* virgins. Mutant virgins behave, and respond to neuronal manipulations, in the same way as mated females. Shaded region corresponds to AbG in wildtype (grey) and *mir-iab-4/8* knockouts (blue). Mann-Whitney non parametric test, ** p<0.01, *** p<0.001, **** p<0.0001. Error bars= SEM. Scale bar= 25μm. *mir-iab-4/8−/−* mutants=*mir[*Δ*/C11]* transheterozygotes.

Despite their prominent expression in the VNC, where the BX-C miRNAs are selectively deployed, some of these Gal4 lines have certain expression in the brain (Feng et al., 2014). We sought to distinguish the contributions of VNC or brain neurons to the phenotypes of *mir-iab-4/8* null females. To do so, we exploited an *Otd-nls:FLPo* transgene that is expressed in the central brain (Asahina et al., 2014). By using this in combination with FRT-cassettes that excise tub-Gal80 or activate Gal4, we were able to restrict *NaChBac* activation to either the brain or the VNC of miRNA mutant animals (**Figure 3D**). For these tests, we employed the *VT-7068* driver, which showed the most robust rescues (**Figure 3C**). While activation of *VT-7068*-brain neurons did not affect virgin egg-laying by *mir-iab-4/8* mutants, specific activation of *VT-7068*-VNC neurons was sufficient to rescue this mutant defect (**Figure 3C**). Moreover, hyperactivation of only the four VNC ascending neurons labeled by *SAG-1-Gal4*, which has no reported expression in the brain (Feng et al., 2014), partially reduced egg-laying in mutants (**Figure 3C**), although not to the extent seen with *VT-7068*.

Together, these experiments suggest that *mir-iab-4/8* loss compromises the output of the ascending post-mating circuit of virgin females in the VNC, partially evoking the effects of SP signaling after mating (**Figure 3D**).

### Virgin behaviors are sensitive to *hth* levels in switch neurons

miRNAs are generally considered to exert their function by repressing broad target networks, although most regulatory interactions are mild (Bartel, 2018). At the same time, the majority of miRNA knockouts in diverse species have at best subtle phenotypes in development or physiology (Lai, 2015). This has led to the concept that miRNAs collectively regulate multiple transcripts to buffer transcriptional noise, and that individual phenocritical targets may be rare, notwithstanding the fact that miRNAs were first identified on account of specific targets that drive miRNA mutant defects (Lee et al., 1993; Wightman et al., 1993).

In previous studies, we identified a network of Hox genes and their TALE-class cofactors as direct *in vivo* targets that are misexpressed in *mir-iab-4/8* mutants, and exhibit dominant phenotypic interactions with the miRNA deletion (Garaulet et al., 2014; Garaulet and Lai, 2015). We were particularly interested in the post-transcriptional regulation of *homothorax* (*hth*), which exhibits the highest number of miRNA binding sites for all the BX-C miRNAs, most of them encoded in a 2.3kb 3’ UTR extension found only in *hth* neural transcripts (Garaulet et al., 2014). Hth protein accumulates broadly across the thoracic neurons of the larval VNC, but is largely excluded from abdominal segments where it overlaps BX-C miRNAs. The BX-C miRNA mutants derepress Hth in abdominal neurons (**Figure 4A**, **Supplementary Figure 5A**) (Garaulet et al., 2014). Since we documented behavioral defects in adults, we also analyzed *hth* patterning in the CNS of adult females. By co-labeling Hth with the tub-GFP-miR-iab-4 sensor and Abd-A in wildtype, we can register that Hth is normally largely excluded from the abdominal ganglion region that encompasses active miR-iab-4 (**Figure 4C, D, G** and **Supplementary Figure 5C**). By extension, Hth is also mostly excluded from the miR-iab-8 domain that is located more posteriorly (**Figure 4C, D, G** and **Supplementary Figure 5C**). As in larval VNC (**Figure 4B** and **Supplementary Figure 5A**), we observed broadly ectopic Hth that extends into miR-iab-4/8 domains in *mir-iab-4/8* mutants, throughout the dorsal and ventral regions of the abdominal ganglion both in female (**Figure 4E, F, H**), and male VNCs (**Supplementary Figure 6D**). Importantly, although there is heterogeneous distribution of nuclei with high levels of ectopic Hth (**Supplementary Figure 6A**), we consistently observe a lower level of derepressed Hth throughout the entire abdominal ganglion (**Figure 4J, J’** and **Supplementary Figure 5B**). This region encompasses populations of neurons that are functionally relevant to the post-mating switch (Feng et al., 2014), including neurons defined by the *VT-7068*, *VT-50405* and *SAG-1* drivers (**Figure 4I-J** and **Supplementary Figure 6B,C**). By contrast, we found no evidence for misexpression of Hth in mutant female brains, where miR-iab-4/8 are not expressed (**Figure 4K-L’**).

**Figure 4.**
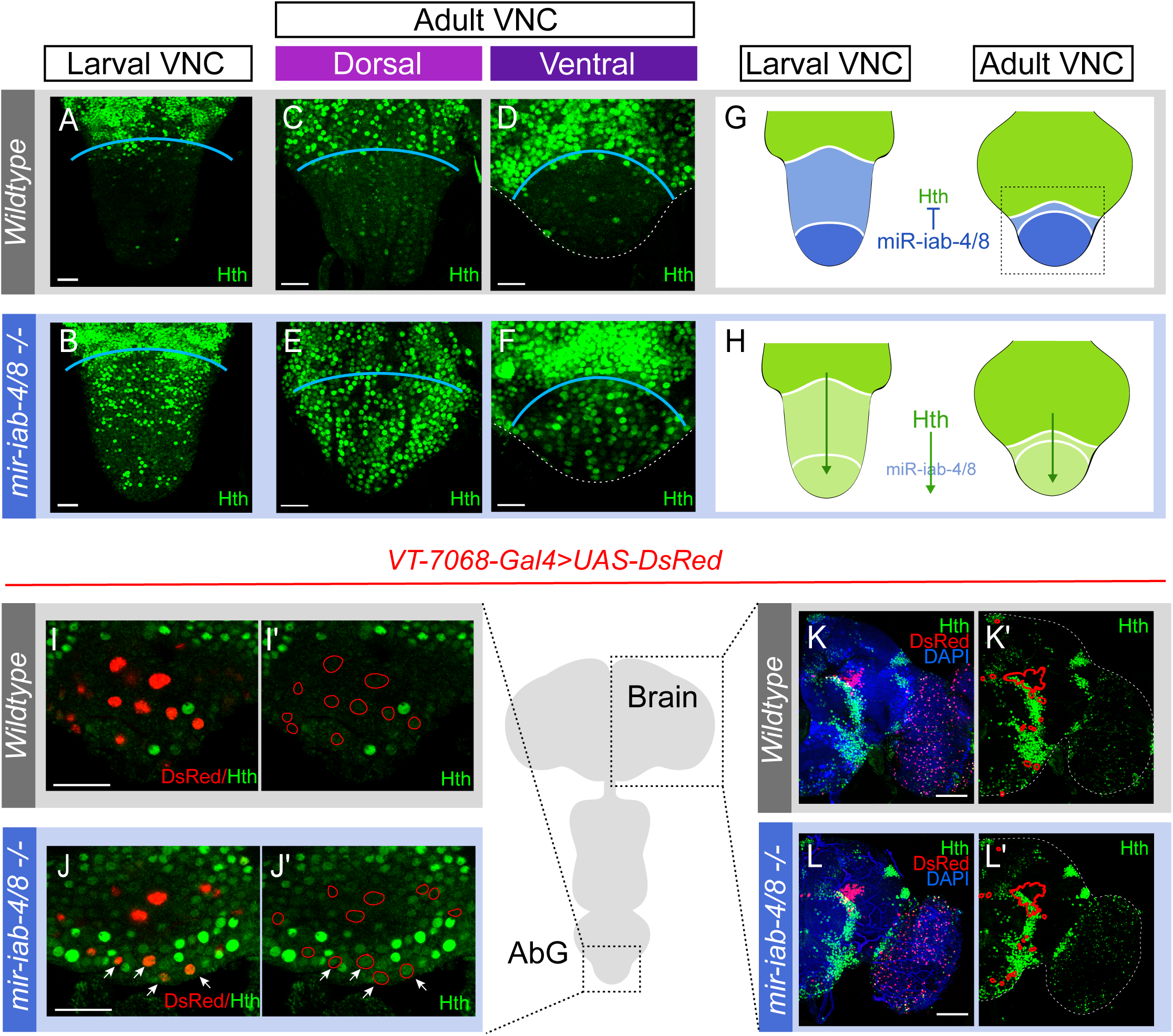
Hth patterning in wildtype and *mir-iab-4/8* mutant CNS. (A-F) Confocal images of female AbGs (boxed region in G). Hth (green) is mostly excluded from the abdominal segments in wildtype females in larval (A) and adult VNC (dorsal in C, ventral in D). Nevertheless, it is widely accumulated *mir-iab-4/8* mutants (B, E, F). (G,H) Schematic representation of abdominal patterning of *hth* by BX-C miRNAs in wildtype (G) and *mir-iab-4/8* mutants (H). *hth* is not expressed in VT-switch neurons in wildtype VNCs (I-I’) but detectable in *mir[*Δ*/C11]* mutants (arrows, J-J’). Hth is expressed in discrete clusters in the female brain, with restricted overlap with VT-7068 neurons (K,K’). Unlike abdominal neurons, no *hth* misexpression is observed in the brain (L,L’). Scale bar= 25μm (A-F, I-J’) and 60μm (K-L’).

We used genetic strategies to evaluate if misexpression of Hth in those specific populations of neurons is responsible for the behavioral shift in *mir-iab-4/8* null virgins. By monitoring egg-laying, we did not observe dominant rescue of the mutant phenotype by heterozygosity of *hth[P2]* null allele, or by a deficiency that uncovers *hth* (**Figure 5A**). However, we observed substantial rescue of miRNA mutant virgin egg-laying upon *hth* knockdown in several sets of VT-switch lines, particularly with *VT-7068* and *VT-50405* (**Figure 5B**). Suppression of *hth* in the intersection of these two Gal4s (*SAG-1* neurons) also rescued partially, indicating involvement of these specific cells (**Figure 5B**). However, the weaker effect suggests that derepression of Hth in non-overlapping sets of *VT-7068* and *VT-50405* neurons also contributes to the miRNA phenotype. Amongst VT-switch lines tested, the strongest Hth accumulation was found in some of the *VT-454* neurons (**Supplementary Figure 6C**). Interestingly, while *VT-454>hth[RNAi]* alone did not suppress egg-laying, inclusion of a *hth* mutant allele combined to yield nearly complete rescue (**Figure 5C**). This genetic threshold effect, not seen with *hth* heterozygosity alone, further highlights the sensitivity of this behavioral defect to Hth levels.

**Figure 5.**
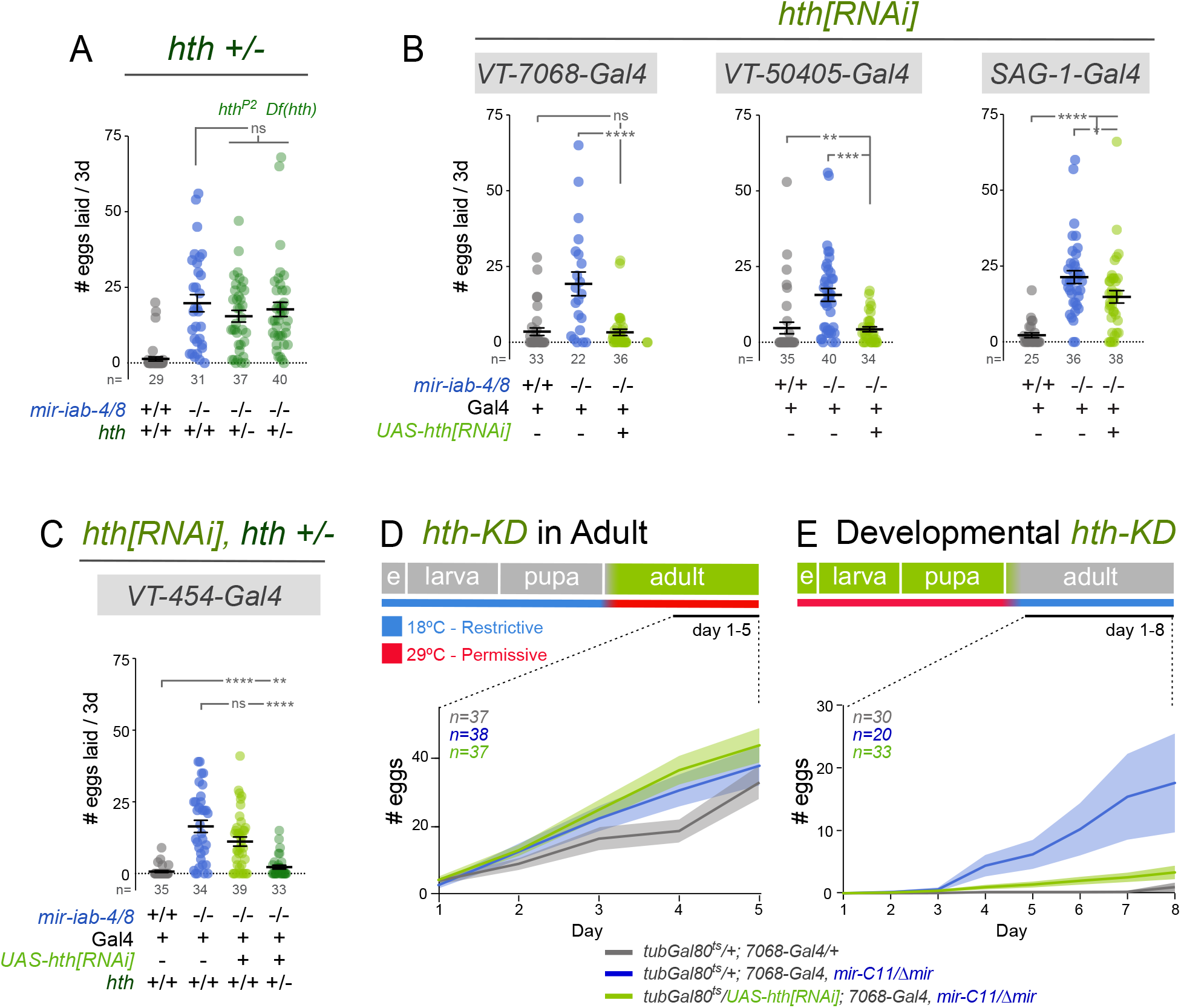
Developmental derepression of Homothorax in VT-switch neurons compromises virgin behavior. (A-C) *hth* dosage manipulations. (A) Heterozygosity of *hth* is not sufficient to modulate increased egg-laying in miRNA mutant animals. (B) Conversely, *hth* knockdown in specific sets of neurons in *mir-iab-4/8* mutants is sufficient to completely (*VT-7068* neurons) or partially (*VT-50405* and *SAG-1* neurons) revert egg-laying to wildtype virgin values. (C) With *VT-454*, neural *hth* knockdown and heterozygosity act synergistically to rescue egg-laying. (D,E) Developmental vs. adult requirements for *hth* regulation by BX-C miRNAs in the PMR. Adult specific knockdown of *hth* in *VT-7068* neurons does not revert elevated egg-laying observed in *mir-iab-4/8* mutants, despite the higher baseline levels observed for wildtype flies in this temperature plan (D). Conversely, developmental knockdown almost completely rescued mutant egg-laying to wildtype values (E). Mann-Whitney non parametric test, ns=not significant, * p<0.05, ** p<0.01, *** p<0.001, **** p<0.0001.. Error bars= SEM. Shaded area (D,F)= SEM. Wildtype flies are *Canton-S* strain, *mir-iab-4/8 −/−* mutants=*mir[*Δ*/C11]* transheterozygotes.

Although *7068-Gal4* drives specific expression in the brain (**Figure 4K-L**), VNC specific rescue of *hth* knockdown using this line was sufficient to substantially revert egg-laying in *mir-iab-4/8* females (**Supplementary Figure 7**), highlighting the relevance of *hth* regulation of specific sets of abdominal neurons where miR-iab-4/8 are expressed.

Since PMRs were triggered by Hth misexpression in *mir-iab-4/8* mutants, we wondered if behavior changes in mated females might involve acute changes in Hth levels in relevant abdominal neurons following copulation. However, immunostaining of the VNCs of wildtype females did not reveal dynamic elevation of Hth in abdominal segments after mating (**Supplementary Figure 6E**). This suggests that upregulation of Hth interferes with the proper function of mutant switch neurons, but is not involved in the normal response to copulation.

Hth upregulation in the VNC is observed throughout development and adult life. To discern the temporal requirements of miR-iab-4/8 regulation of *hth*, we restricted *hth* knockdown to either developmental stages or adult life, by inclusion of *tubGal80^ts^* transgene, and placing the flies at restrictive (18°C) or permissive temperature (29°C) before or after eclosion. (**Figure 5D-E**). Of note, raising the temperature after eclosion stimulates egg-laying in adult wildtype females (**Figure 5D** compare to **Figures 1A**, **3A** and **5A**). Nonetheless, *mir-iab-4/8* mutant females still lay comparatively more eggs under these temperature conditions, and adult specific *hth* knockdown after eclosion does not revert this trend (**Figure 5D**). By contrast, temperature elevation during development did not affect egg-laying in wildtype virgins (**Figure 5E**). In this setting, developmental silencing of *hth* in *VT-7068* neurons was sufficient to revert the elevated oviposition rates observed in *mir-iab-4/8* nulls (**Figure 5E**).

Overall, while Hth is broadly misexpressed throughout the miR-iab-4/8 domains in mutants, its post-transcriptional repression in restricted sets of abdominal neurons during development mediates the capacity of virgins to integrate internal state and behavior.

### Direct evidence that BX-C miRNA binding sites in the *hth* 3’ UTR control virgin behavior

Genetic interactions can support compelling linkages amongst pathway components, but do not alone prove direct regulation. Thus, there is growing impetus to utilize genome engineering to specifically interrogate specific miRNA-target interactions *in vivo* (Ecsedi et al., 2015). While this is more straightforward for loci with one or two sites of interest, the full-length homeodomain-encoding isoform *hth-HD* bears 15 conserved sites for miR-iab-4/8, (two of which are predicted as dual sites). In order to disrupt all of these sites at the endogenous locus, we implemented a multi-step strategy that permits flexible engineering (**Figure 6A**, **Supplementary Figure 8A**). We first used CRISPR/Cas9 to replace the last two exons and 3’ UTR with an attP site (**Figure 6B**, **Supplementary Figure 8A**). The resulting *hth-HD[*Δ*attP]* allele was homozygous lethal, in accordance with known deletions of the distal end of *hth* isoforms (Kurant et al., 1998; Pai et al., 1998). This engineerable allele is amenable to generation of diverse downstream variants using ΦC31-mediated insertion (Baena-Lopez et al., 2013) (**Figure 5C**).

**Figure 6.**
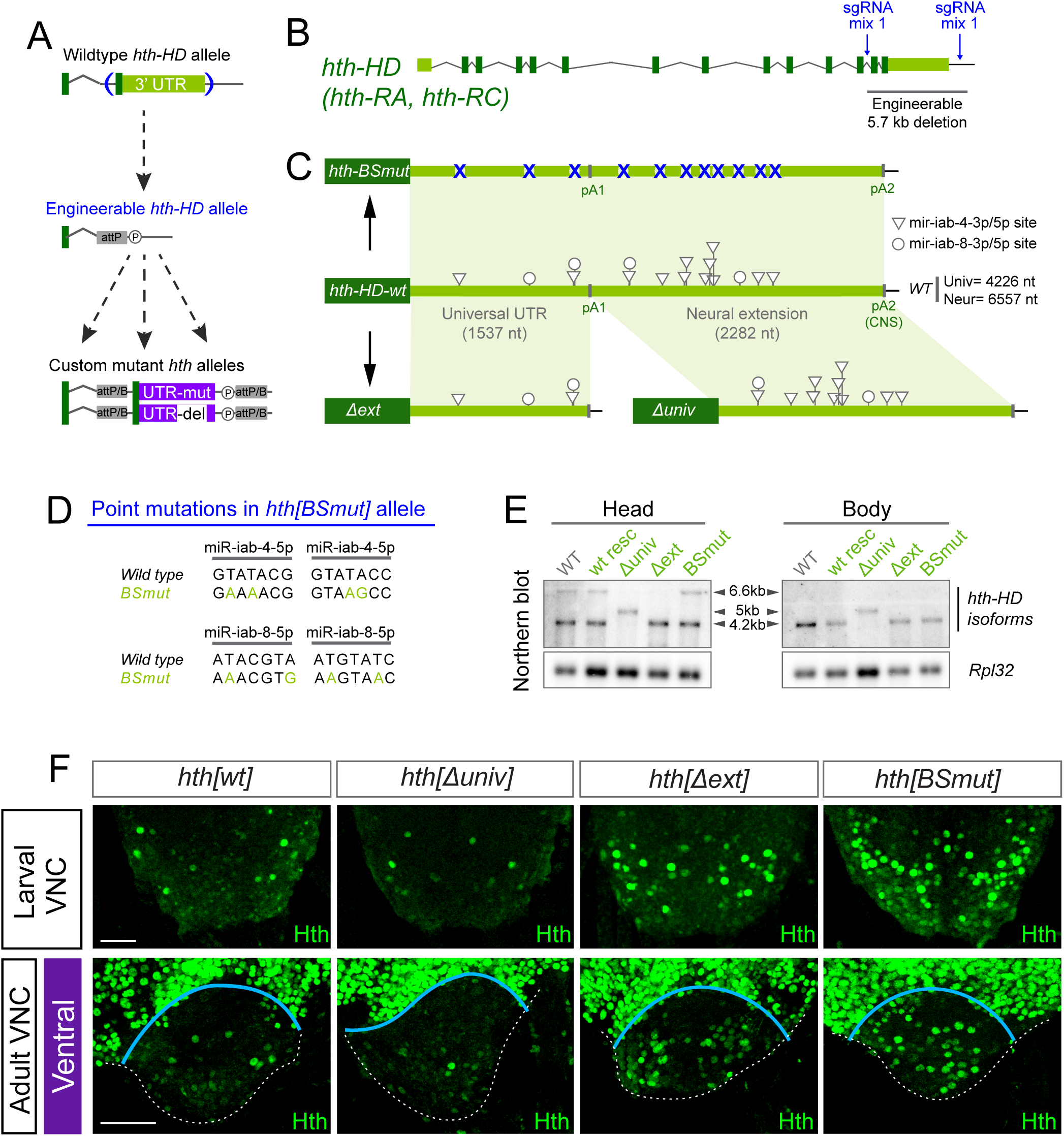
Engineering of *homothorax* 3’UTR. (A) Flexible 3’ UTR engineering pipeline. (B) sgRNA cocktails were designed to delete the last 2 exons and the full 3’UTR of *hth-HD* isoform, which encodes the full-length homeodomain (HD) factor. (C) *hth-HD* 3’UTR alleles generated by CRISPR/ΦC31 mediated deletion/replacement. (D) Two point mutations were introduced per predicted binding site for either mature species of miR-iab-4 and miR-iab-8. (E) Northern blotting of the *hth-HD* engineered alleles. Deletion of the neural extension (*hth[*Δ*ext]*) eliminates the 6.6 kb isoform, only present in heads. Deletion of the universal segment (*hth[*Δ*univ]*) generates a novel isoform of 5kb, which constitutively incorporates the UTR extension both in heads and bodies (5kb). *hth[wt]* replacement and binding sites mutant (*hth[BSmut]*) do not affect the isoforms detected. kb=kilobase. (F) Hth pattern in the posterior region of the larval VNC and the AbG of *hth-HD* 3’ UTR alleles. *hth[BSmut]* and *hth[*Δ*ext]* fail to repress *hth* expression in abdominal segments. Scale bar= 50 μm.

Here, we re-introduced either the deleted segment of the native gene (wildtype rescue, *hth-[wt]*), or a 3’ UTR variant in which two seed bases of all conserved miR-iab-4 and miR-iab-8 binding sites were mutated (*hth-[BSmut],* **Figure 6C-D**). Thus, we obtained directly comparable control and modified *hth* alleles with only minimal genomic scars. Northern blotting confirmed that these replacement alleles preserved the biogenesis of proximal and distal *hth* isoforms of expected sizes (**Figure 6E**). In both cases, full viability of *hth-HD[*Δ*attP]* was restored, and no external developmental defects were observed. By contrast, immunostaining of the adult VNC revealed substantial defects specifically in the *hth[BSmut]* replacement allele. In particular, Hth was derepressed in the mutant abdominal ganglion compared to the wildtype 3’ UTR replacement, both during larval stages (**Supplementary Figure 8B**) and in adult females (**Figure 6F** **and Supplementary Figure 8C**). The adult de-repression of Hth was more overt in the ventral VNC relative to dorsal VNC (**Supplementary Figure 8C**). Overall, these experiments yield stringent evidence that specific endogenous miRNA sites are essential to exclude Hth from the abdominal ganglion (**Figure 4G, H**).

Since the spatial derepression of Hth was less in *hth[BSmut]* (**Figure 6F** **and Supplementary Figure 8C**) than with deletion of the miRNAs *per se* (**Figure 4E-F**), it was reasonable to wonder if this is sufficient to drive behavioral changes. Strikingly, we found that *hth[BSmut]* females nearly phenocopy the switch to the mated state observed in miRNA nulls. Compared to *hth[wt]*, which exhibits normal virgin behaviors, *hth[BSmut*] virgins lay substantial numbers of eggs, exhibit reduced receptivity to courtship, fail to open their vaginal plates, but perform ovipositor extrusions instead (**Figure 7A-D**). Thus, in spite of other evident misregulation of Hth that occurs in the miRNA deletion, perhaps reflecting indirect effects involving other Hox gene targets (Garaulet et al., 2014), failure of its direct repression by miR-iab-4/8 can account for the uncoupling of internal state and behavioral output in virgin females with respect to multiple PMR readouts (**Figure 6E** **and Supplementary Figure 9**).

**Figure 7.**
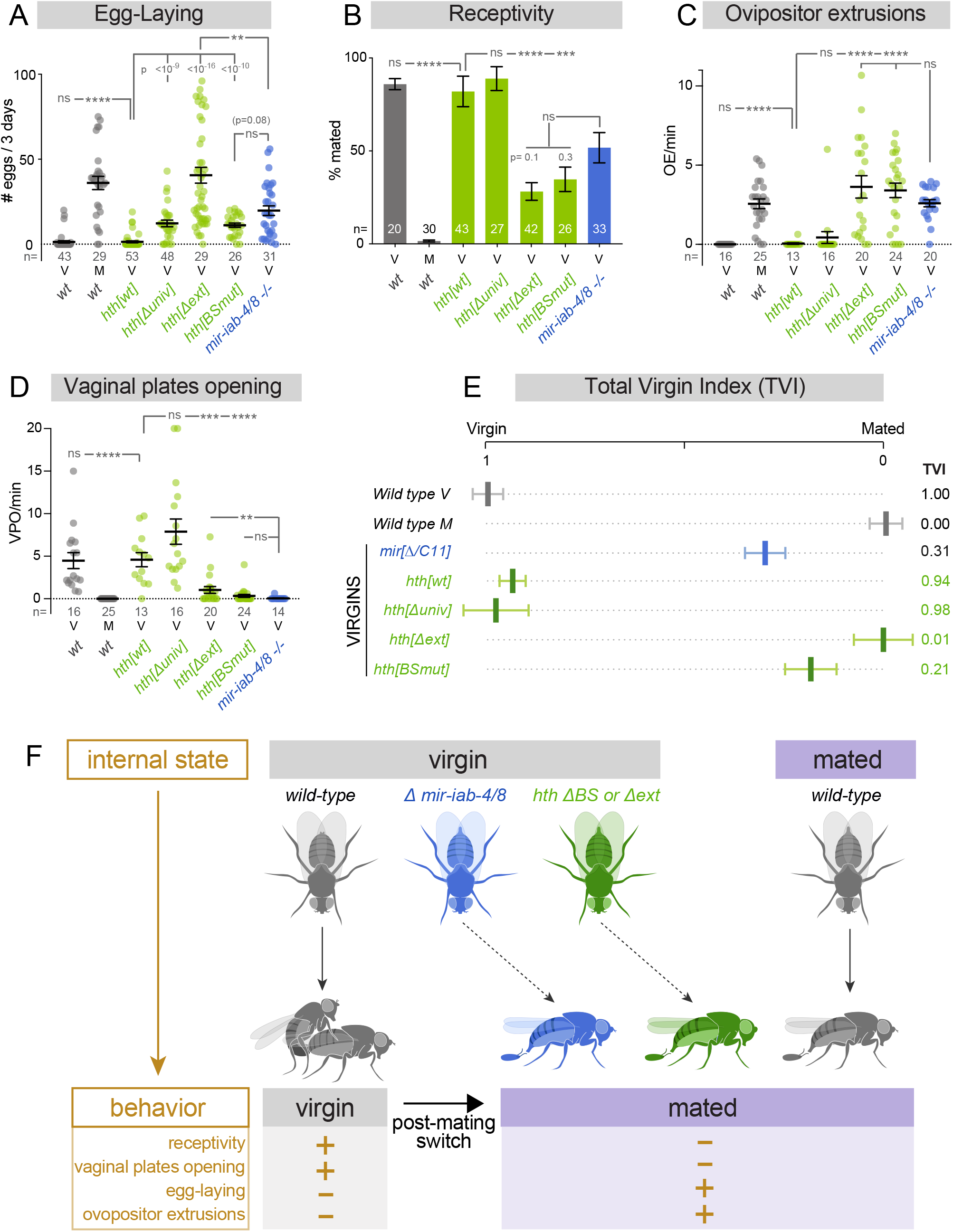
Functional study of *hth-HD* 3’UTR elements. (A-D) Virgin and mated behaviors assayed in wildtype, *mir-iab-4/8* and *hth-HD* 3’ UTR alleles. (A) Egg-laying, (B) receptivity, (C) ovipositor extrusions, and (D) vaginal plates openings. Deletion *hth-HD* 3’ UTR neural extension or mutation of miR-iab-4/8 binding sites inhibit virgin behaviors and induce mated behaviors in virgins. (E) Total virgin index, integrating the values of all four behaviors analyzed in A-D. (F) Summary illustrating how neural specific post-transcriptional regulation of *homothorax* is required for appropriate female behavior. Multiple behaviors in virgin females are altered after mating. The mated state is induced in virgin females deleted for *mir-iab-4/8*, by specific mutation of their binding sites in endogenous *hth*, or by deletion of the *hth* neural 3’ UTR extension. Thus, the integration of both miRNA control and neural APA is required to execute behaviors that are appropriate to the internal state of virgin female flies. (B) Fisher’s exact test. (A, C, D) Kruskal-Wallis t test with Dunn’s correction. ns=not significant, * p<0.05, ** p<0.01, *** p<0.001, **** p<0.0001. Error bars= SEM. Wildtype flies are *Canton-S* strain, *mir-iab-4/8−/−* mutants= *mir[*Δ*/C11]* transheterozygotes.

### Integration of neural APA with miRNA regulation adjusts virgin behavior to internal state

Neural 3’ UTR extensions induced by tissue-specific polyadenylation adds substantial regulatory real estate to hundreds of transcript isoforms (Sanfilippo et al., 2017). However, there are few *in vivo* examples of such extensions controlling phenotypically overt processes (An et al., 2008; Andreassi et al., 2010; Zhang et al., 2019). There are conserved miR-iab-4 and miR-iab-8 sites in both the universal and extended 3’ UTR segments of *hth* (Garaulet et al., 2014), although more total sites reside in the neural extension (**Figure 6C**). We prepared two additional *hth* alleles, in which we deleted either the universal or extended regions of the *hth-HD* 3’ UTR (**Figure 6C**). We confirmed that these alleles yielded expected transcript sizes, that is, deletion of the universal portion (Δ*univ*) resulted in a single common isoform bearing the extended 3’ UTR in head and body RNA, whereas deletion of the extension region (Δ*ext*) yielded a single short isoform in both RNA samples (**Figure 6E**).

Notably, these alleles had disparate effects on Hth protein expression and animal behavior. In particular, *hth[*Δ*ext]* caused far more abdominal neurons to accumulate Hth than did the *hth[*Δ*univ]* allele (**Figure 6F and Supplementary Figure B,C**). Because the universal *hth* 3’ UTR segment contains multiple conserved miR-iab-4/8 sites, it is conceivable that loss of this regulatory region impacts VNC development. We found that virgin *hth[*Δ*univ]* mutants exhibited elevated egg-laying, but otherwise had normal sexual receptivity and genital behaviors. Thus, virgin behaviors seemed mostly intact in *hth[*Δ*univ]* mutants. By contras, *hth[*Δ*ext]* virgins displayed full PMRs with respect to all behavioral parameters, namely egg-laying, sexual receptivity, ovipositor extrusions, and frequency of vaginal plate openings (**Figure 7A-D** and **Supplementary Figure 9**).

We integrated their performances in multiple assays into a single value, total virgin index (TVI). We assigned equal contributions to the different PMR readouts, which we acknowledge is arbitrary, but provides a simple metric to aid in the comparison of overall behaviors across mutants. This comparison emphasizes that the specific *hth* engineered mutants cause stronger induction of mated responses in virgins than does deletion of the miRNA locus itself (**Figure 7F** **and Supplementary Figure 9**). In fact, *hth[*Δ*ext]* virgins are nearly indistinguishable from wildtype mated females. Since deletion of the miRNA locus may alter other direct and potentially indirect targets (Garaulet et al., 2014), we may not expect a complete correspondence of phenotypes, although these are qualitatively similar. On the other hand, as all the *hth* mutants are controlled with respect to a precise replacement of the wildtype 3’ UTR, these data highlight the strength of precision genetics to assay *in vivo* requirements at an individual phenocritical locus. Overall, these data demonstrate that miRNA regulation and neural APA are integrated at the *hth* locus to enable the virgin behavioral state in *Drosophila* females.

## Discussion

### Intersection of miRNA control and neural alternative polyadenylation

miRNAs are known to influence diverse aspects of development and adult physiology, and some miRNA mutants have been suggested to affect post-mated responses (Bender, 2008; Fricke et al., 2014; Garaulet et al., 2014; Maeda et al., 2018). Nevertheless, many miRNA knockout effects are subtle and challenging to comprehend using classical genetics and epistatic analysis. These strategies were critical to initially uncover the biological impact of miRNAs and to determine key individual miRNA targets whose regulation/dysregulation mediates *in vivo* phenotypes (Lai et al., 1998; Lee et al., 1993; Leviten et al., 1997; Wightman et al., 1993). Although we have learned much about the quantitative influence of miRNA targeting across the transcriptome (Agarwal et al., 2015; Kim et al., 2016), computational and genomic strategies still cannot predict which miRNAs and targets are involved in phenotypically overt regulation, and what settings these might manifest in. The finding that miRNA targeting is highly conditional on 3’ UTR utilization, via alternative polyadenylation (APA), raises further complexities into considering miRNA biology (Gruber and Zavolan, 2019). Nevertheless, it has also been reported that at least in some contexts, such as cultured fibroblasts, APA has limited impacts on gene expression and translation (Spies et al., 2013).

In this study, we dissect the biological impact of miRNAs and APA, which intersect to regulate an individual critical target in the nervous system. In particular, we use genetic engineering and neurobiological strategies to reveal a biological imperative for alternative polyadenylation that extends the *hth* 3’ UTR in the CNS. In general, many hundreds of genes express extended 3’ UTRs in the nervous system, which presumably increase post-transcriptional capacities of these neural isoforms (Miura et al., 2014). However, there are few *in vivo* reports that such neural 3’ UTR APA extensions mediate biologically substantial effects (An et al., 2008; Terenzio et al., 2018). By taking advantage of our engineerable allele platform, we were able to delete the universal and extended *hth* 3’ UTR in the intact fly. These mutants provide evidence that the role of neural APA is to unveil an array of BX-C miRNA sites that are crucial for *hth* suppression in the posterior VNC, where the miRNAs are normally expressed. Since BX-C miRNAs do not appear to be active in other settings where Hth is involved in patterning, e.g., in imaginal discs, the APA mechanism allows for heightened sensitivity of Hth to miRNAs in the CNS. Other BX-C targets of the BX-C miRNAs similarly undergo neural APA (Garaulet et al., 2014; Thomsen et al., 2010). Although phenotypic demonstration of their regulatory impacts of these extensions remain to be elucidated, the engineering platform described here is suitable. Moreover, it will be of considerable interest to determine the mechanism by which neural APA extensions of these and other genes are achieved (Hilgers et al., 2012).

### Genetic control of the virgin behavioral state

We exploited the entry point of a miRNA knockout, which unexpectedly bypasses the virgin behavioral state of females. Multiple loss-of-function mutants are known (e.g., of SP and SP receptor) in which females cannot transition to normal mated behaviors following copulation. However, while certain transgenic interventions can induce mated conducts in virgins, such as inhibiting PKA activity downstream of the SP receptor in SPSNs (Yang et al., 2009), deletion of the BX-C miRNAs appears to be the first loss-of-function mutant that induces the mated state in virgins. Moreover, this effect does not appear to be the consequence of broad derepression of a large target cohort, but instead can be largely attributed to a single key target (*hth*).

These findings have important implications for understanding genetic impacts on the post-mating switch. While it has been frequently assumed that the virgin state is a default state, our results reveal genetic programming of virgin behavior. We use precision genetics to identify two new factors that are critical for the execution of virgin behaviors. Moreover, we reveal how the neural exclusive 3’ UTR extension of *hth* installs critical post-transcriptional elements, principally BX-C miRNA sites, which restrict Hth expression in the CNS and mediate appropriate adult behaviors. We observe derepression of the causal target Hth during both larval and in adult stages, but our spatio-temporally controlled experiments demonstrate that misexpression of Hth in the developing VNC is critical for the defective post-mating response. We note that the strong misexpression of Hth in adult VNC neurons of miRNA mutants likely has consequences for gene regulation, directed misexpression of this transcription factor has powerful detrimental consequences. However, it remains to be seen how its adult-specific misexpression affects behavior, since the phenotypes we analyzed here appear to have mostly developmental origins.

In any case, it is clear that both BX-C miRNAs and Hth comprise an unanticipated developmental regulatory axis that is required for adult virgin behavior. The failure of appropriate Hth regulation has two consequences for female virgin flies. First, they do not readily copulate, although we do observe that they can be inseminated. However, they also inappropriately execute reproductive activities without having mated. As these behaviors such as egg-laying are energetically demanding and reduce female fitness, it is necessary to avoid these before insemination, when they would be otherwise unsuccessful (**Figure 7F**). We reveal that females encode dual post-transcriptional regulatory mechanisms that are required to execute behaviors that are appropriate to the internal state of virgin flies.

## STAR METHODS

### KEY RESOURCES TABLE

|  | SOURCE | IDENTIFIER |
| --- | --- | --- |
| **Antibodies** |  |  |
| rabbit anti-Hth (1:500) | Salvany et al, 2009 | N/A |
| guinea pig anti-Hth (1:500) | Salvany et al, 2009 | N/A |
| Mouse anti-Abd-A (1:500) | Gift of Ian Duncan | N/A |
| **Bacterial and Virus Strains** |  |  |
| TOP10 Chemically Competent E. coli | Our lab | N/A |
| **Chemicals, Peptides, and Recombinant Proteins** |  |  |
| Penicillin-Streptomycin | Life Technologies | Cat#15070063 |
| T4 DNA Ligase | NEB | Cat#M0202L |
| Gel Loading Buffer II | Invitrogen | Cat#AM8547 |
| AccuPrime pfx DNA Polymerase | Thermo Fisher | Cat#12344024 |
| Trizol reagent | Life Technologies | Cat#15596018 |
| **Experimental Models: Organisms/Strains** |  |  |
| mir[Δ] | gift of Welcome Bender (Bender, 2008) |  |
| Canton-S | gift of Karla Kaun |  |
| VT-lines and SAG-1 split Gal4 line | gift of Barry Dickson (Feng et al., 2014) |  |
| UAS-hth-RNAi, | Vienna Drosophila RNAi Center |  |
| hth[P2] | gift of Richard Mann (Pai et al., 1998) |  |
| Df[hth], | Exelixis Collection |  |
| Otd-nls:FLPo, | gift of David J. Anderson |  |
| tub-GFP-sensors for miR-iab-4 and miR-iab-8 | (Garaulet et al., 2014) |  |
| Oregon R | Bloomington Drosophila Stock Center |  |
| Crimea | Bloomington Drosophila Stock Center |  |
| UAS-Red-Stinger | Bloomington Drosophila Stock Center |  |
| nos-cas9 (attP40) | Bloomington Drosophila Stock Center |  |
| nos-ΦC31 (X) | Bloomington Drosophila Stock Center | |
| TM6B-hs-Cre, | Bloomington Drosophila Stock Center |  |
| >tubGal80> | Bloomington Drosophila Stock Center |  |
| tub>stop>Gal80 | Bloomington Drosophila Stock Center |  |
| tubGal80ts | Bloomington Drosophila Stock Center |  |

### CONTACT FOR REAGENT AND RESOURCE SHARING

Further information and requests for resources and reagents should be directed to and will be fulfilled by the Lead Contact, Eric Lai, or by Daniel L. Garaulet.

### EXPERIMENTAL MODEL AND SUBJECT DETAILS

Wildtype and genetically engineered strains of *Drosophila melanogaster* were used in the study. Larval and adult flies were raised on standard molasses fly food and kept at 25°C, 55% humidity unless mentioned otherwise.

## METHOD DETAILS

### Drosophila stocks

*mir[*Δ], gift of Welcome Bender (Bender, 2008)

Canton-S, gift of Karla Kaun

VT-lines and SAG-1 split Gal4 line, gift of Barry Dickson (Feng et al., 2014)

*UAS-hth-RNAi*, Vienna Drosophila RNAi Center

*hth[P2]*, gift of Richard Mann (Pai et al., 1998)

*Df[hth]*, Exelixis Collection

*Otd-nls:FLPo*, gift of David J. Anderson

*tub-GFP*-sensors for miR-iab-4 and miR-iab-8 (Garaulet et al., 2014)

The following lines were obtained from Bloomington Drosophila Stock Center: *Oregon R* and *Crimea* strains, *UAS-Red-Stinger*, *nos-cas9* (attP40), *nos-*Φ*C31* (*X*), *TM6B-hs-Cre, >tubGal80>, tub>stop>Gal80, tubGal80^ts^*. All the lines used in this study have been backcrossed at least 8 generations to the *Canton S* wildtype strain.

### CRISPR-Cas9 engineered alleles

*mir-iab-4/8* deletion [*C11*] was generated using a CRISPR transgenic approach (Kondo and Ueda, 2013). The sgRNA guide: ATACTGAAGGTATACCGGAT-AGG (chr3R:12682031-12682045) was inserted under the U6 promoter in Chr III, and subsequently crossed to *nos-cas9* expressing flies (BDRC 78781) to mutagenize *mir-iab-4/8* locus. Individual offspring were analyzed by PCR and sequencing to detect mutant events. The potential structures of miRNA deletion transcripts were predicted using RNAfold (http://rna.tbi.univie.ac.at/cgi-bin/RNAWebSuite/RNAfold.cgi). *hth-HD* deletion was generated by direct injection of a CRISPR mix containing multiple pCFD4 dual sgRNAs at 75ng/µL each (Port et al., 2014), and a single pHD-attP-DsRed donor at 100 ng/µL (Gratz et al., 2014) in nos-Cas9 expressing flies (BDRC 78781).

### sgRNA sequences

Intron break:

GAGGCCGTGTCAAAGCTTGC-GGG chr3R: 6337918-6337932

GTGACTGTCACTCGGCCCGC-CGG chr3R: 6337877-6337891

TTAAAGCAGAAAGCCGCCGG-CGG chr3R: 6337869-6337883

GCAGTTGCCCGTTGATCCGC-AGG chr3R: 6337807-6337821

GCTTGCGGGTAACATCGTCT-TGG chr3R: 6337904-6337926

### Downstream break

CGAGCTGGTGGCTCTCCCGC-TGG chr3R: 6332418-6332432

AAAGTAATTACAGCCGCCGA-CGG chr3R: 6332151-6332165

TGCCTATAATTAGTGGGCAG-TGG chr3R: 6332216-6332230

### Primers used to clone homology arms in pHD-att-DsRed

Intronic Homology Arm:

Fwd; AATATCGCATCGCCCCC chr3R: 6338982-6339004

Rev; AGCTTTGACACGGCCTC chr3R: 6337921-6337940

Fwd; GCGGCTGTAATTACTTTTACG chr3R: 6332139-6332159

Rev: AATAAATGGCTTGACAAACGC chr3R: 6330834-6330854

Mutant flies were identified by fluorescent red eyes. Appropriate substitution events were characterized by PCR and sequencing. Reintegration of custom UTR sequences was performed using pRIV-attB vectors(Baena-Lopez et al., 2013) in nos-ΦC31 expressing flies (DBSC 34771). *hth[BSmut]* was obtained by stitching together multiple g-block – Gene Fragments (Integrated Gene Technologies) carrying point mutations in conserved miR-iab-4/8 binding sites. Injections were performed at Bestgene, Inc. and Rainbow Transgenic Flies, Inc. All new alleles were isogenized to the Canton-S strain before behavior analysis.

### Northern Blotting

Body and head samples were prepared using liquid nitrogen to freeze, followed by vortexing and separation of heads from bodies using a sieve. Total RNAs were extracted from either heads or bodies of all genotypes using TRIzol (Invitrogen). Poly(A)+ RNAs were enriched from total RNAs using Oligo d(T)25 magnetic beads (New England Biolabs), according to the manufacturer’ s instructions. Northern blot analysis was performed essentially as previously described (Smibert et al., 2012) using 1.5 μg of poly(A)+ RNA per lane. Millennium RNA Marker (Invitrogen) was used as RNA size standard for each blot. Northern blot probes to detect different isoforms of homothorax were prepared by Amersham Megaprime DNA Labeling System (GE Healthcare Life Sciences), according to manufacturer’s instructions. Oligonucleotide sequences used to generate northern probes are listed in Table S1.

### Behavioral assays

Virgin males and females were collected after eclosion and were kept isolated in vials at 25°C, 55% humidity and 12h:12h LD cycles until the time of the behavior test. Virgin behaviors were assayed 3 days after eclosion unless specifically mentioned otherwise. For the analysis of mated females, they were mated a first time at day 3, separated from male after copulation, and kept isolated for 24h until the performance of the mated responses.

Receptivity, ovipositor extrusions and vaginal plates openings were analyzed from video recordings of custom made, 18x multiplex mating arenas (chamber size: 10mm diameter). Single males and females were placed in a half of each arena and let accommodate for 5 min before the assay. Egg-laying was calculated as the number of eggs laid in the first 3 days after eclosion. For receptivity; courtship initiation, mating start, and mating end were manually annotated. Receptivity was calculated as the proportion of animals mating at each time during the course of an hour (Figure 1C), or cumulative proportion of animals mated at 10 min (rest of the figures). Ovipositor extrusions and vaginal plates openings were examined during the first 4 minutes after courtship copulation or until mating. Counts of either behavior were normalized to time. For ovipositor extrusions of *mir-iab-4/8* mutants we used 1 day old virgins, since quantitative levels of extrusions were slightly higher at this age (**Supplementary Figure 2**). Nonetheless, nearly all flies displayed some levels of extrusions at any age from day 1 to 3. All tests were performed at ZT 7-11 and at least at four different occasions. Wildtype and heterozygous control females are always Canton-S strain unless otherwise specified. All males are Canton-S.

Temperature shifts were induced immediately after eclosion. Both at restrictive (18°C) and permissive (29°C) temperature, 55% humidity and 12h:12h LD cycles were maintained.

Virgin Indexes (VI) were calculated by interpolating the behavioral value obtained for each mutant fly within the rank defined from the average wildtype virgin value (1) and the average wildtype mated value (0). The plots show the average and SEM of all the individual values per behavior (Virgin Index), or the average of all behaviors (Total Virgin Index).

Receptivity was analyzed by Fisher’s exact test. Egg-laying, Ovipositor extrusions, vaginal plates opening and cell counts were analyzed by Mann-Whitney non-parametric test or Kruskal-Wallis t test with Dunn’s correction for multiple comparisons.

### Immunohistochemistry, imaging, and image quantification analysis

Larval and adult CNS were dissected in cold 1x PBS and fixed for 1h in 4% paraformaldehyde + 0.1% Triton. Primary and secondary antibodies were incubated for >36h at 4°C in wash buffer (PBS 1% BSA) and mounted in Vectashield (Vector Labs). Antibodies used were rabbit and guinea pig anti-Hth (1:500) (Salvany et al., 2009), mouse anti-Abd-A (1:500), and Alexa-488, −555, −647 conjugated goat and/or donkey antibodies from Thermo Fisher Scientific.

Imaging was performed in a Leica TCS SP5 confocal microscope. Each VNC was typically scanned in 55 planes (Z step ~2nm). When image quantification or comparison was performed (Figures 2,4,6), all different genotypes used were dissected at the same time, fixed and incubated together in the same well. To identify the genotype of each VNC while mounting, different parts of the head were left attached or removed from the VNC during dissection. Then, the same number of VNCs from different genotypes were arranged in a known fashion per slide, to avoid differences in the quantification due to the mounting process. Laser power and offset were maintained identically for all the samples being compared. Gain was slightly adjusted to an internal control in each case. All adult VNC images are 0-24hr old virgin females.

## QUANTIFICATION AND STATISTICAL ANALYSIS

Statistical significance was evaluated using Fisher’s exact test for receptivity (**Figures 1 and 7**), copulation onset (**Figure 1**), qualitative analyses of oviposition and ovipositor extrusions (**Supplementary Figure 1**), and number of neurons (**Figure 2**), and Mann-Whitney non parametric test for Egg-laying, quantitative ovipositor extrusions and vaginal plates openings (**Figures 1,2,3,5,7** and **Supplementary Figures 1,4,7,9**). ns=not significant, * p<0.05, ** p<0.01, *** p<0.001, **** p<0.0001. Error bars in **Figures 1,2,3,4,5,7,9**, and **Supplementary Figures 1,4,7,9**, and shaded area in **Figure 5** represent SEM. All n values are displayed on the figures.

### Materials availability

Plasmid and transgenic flies generated in this study are available from the corresponding authors on request.

## DATA AND SOFTWARE AVAILABILITY

Original behavior data for Figures 1,2,3,5,7 and Supplementary Figures 1,4,7,9 in the paper are available from the corresponding authors on request.

## Acknowledgments

We thank Welcome Bender, Barry Dickson, Natalia Azpiazu, Richard Mann, Carlos Ribeiro, Carolina Rezaval, Stephen Goodwin, Young-Joon Kim, Leslie Vosshall, Shu Kondo, Karla Kaun, Luis Alberto Baena-Lopez, Mariana Wolfner, Ernesto Sánchez-Herrero, Paloma Martin, Michelle Arbeitman, Filip Port, Kate O’Connor-Giles, David J. Anderson and the Bloomington Stock Center for fly strains, plasmids, and antibodies used in this study. Ricardo Toledo-Crow helped fabricate mating chambers. We are grateful to Alex Panzarino and Silvia Rodriguez for helping with initial behavioral assays, and Jennifer Bussell and Leslie Vosshall for advice on assaying female mating behaviors. Work in E.C.L.’s group was supported by the NIH (R01-GM083300 and R01-NS083833) and MSK Core Grant P30-CA008748.

## Author contributions

Conceptualization, D.L.G., E.C.L.; Methodology, D.L.G.; Formal analysis, D.L.G.; Investigation, D.L.G., B.Z.,L.W.,E.C.L.; Resources, D.L.G., E.C.L.; Visualization, D.L.G.; Supervision, D.L.G., E.C.L.; Writing – original Draft and Reviewing and Editing, D.L.G., E.C.L.; Project administration, D.L.G., E.C.L.; Funding acquisition, E.C.L.

## Supplementary Figure Legends

**Supplementary Figure 1 (related to Figure 1). Extended behavioral analysis of *mir-iab-4/8* mutants.** (A) Temporal analysis of sexual receptivity, assessed by the fraction of individual females that are copulating at a given timepoint. The 10 minute timepoint (dotted line) was used subsequently to summarize and compare multiple genotypes or manipulations, shown in Figure 1C. (B) Qualitative oviposition in wildtype and *mir[*Δ*/C11]* flies. Egg-laying is uncoupled from internal state. (C) Quantitative egg-laying counts across several wild type strains. Qualitative (D) and quantitative (E) representation of ovipositor extrusions. Overall levels of male rejection by extruding the ovipositor is maximal in young virgins (1 day old). Fisher’s exact test for qualitative analysis (B, D), and Mann-Whitney non parametric test for quantitative analysis (C, E). ns=not significant, * p<0.05, ** p<0.01, *** p<0.001, **** p<0.0001. Error bars = SEM.

**Supplementary Figure 2 (related to Figure 1).** *mir-iab-4/8* mutant alleles. (A) Wildtype, *mir[*Δ*/*Δ] and *mir[C11/C11]* sequence. (B) RNAfold predictions of each allele. The newly made *mir[C11]* allele deletes miR-iab-8-5p (miR-iab-4-3p) and does not adopt a pre-miRNA-like hairpin structure. Blue= *mir-iab-4/8* species in each allele. Magenta= sgRNA used to mutagenize *mir-iab-4/8 locus*. Low-case letters: inserted nucleotides. nt=nucleotides.

**Supplementary Figure 3 (related to Figure 1). Receptivity dynamics in mir-iab-4/8 mutants.** Time lapse shots from movies of 18 wildtype (*Canton-S*) and 18 *mir[*Δ*/C11]* female flies. In all cases males are Canton-S. Mating couples are marked in yellow. Compared to control, the copulation success of *mir[*Δ*/C11]* mutants is markedly reduced.

**Supplementary Figure 4 (related to Figure 2). Quantification of additional VT-switch neurons.** (A) Schematic of the Drosophila CNS. Boxed region corresponds to the ventral side of AbG shown in B-E. (B) Z-projection of *454-Gal4* VT-switch neurons (cyan) co-stained with Abd-A (red). Number of VT-switch+ cells per VNC domain (*iab-4* vs. *iab-8*). Egg-laying counts in *VT-454>UAS-NaChBac* virgins. Mann-Whitney non parametric test, ** p<0.01, **** p<0.0001. Error bars = SEM. Scalebar = 25μm.

**Supplementary Figure 5 (related to Figure 4). Extended characterization of Hth expression: I.** (A, B) Hth immunostaining (green) in larval VNCs. (A) Hth is derepressed in abdominal segments (Abd-A, red) of *mir[*Δ*/C11]* female larvae (Z-projections). (B) Mild levels of Hth accumulation are detected in nearly all cells of mutant VNCs. Along the dorsoventral axis, Hth is mutually exclusive with miR-iab-4 and miR-iab-8 (inferred from activity GFP-sensors), but not with Abd-A (C). Scalebar= 25μm.

**Supplementary Figure 6 (related to Figure 4). Extended characterization of Hth expression: II.** (A) Hth accumulation in wildtype and two *mir-iab-4/8* mutants, illustrating variability of derepression in mutant VNCs. (B,C) Hth expression relative to VT-switch neuron populations. Only showing the AbG boxed in A. Some VT-454 labeled neurons show especially high accumulation of Hth protein (arrows). (D) Hth expression in AbG of wildtype and *mir-iab-4/8* mutant males is similar to their female counterparts. (E) Copulation does not affect Hth expression levels in the female AbG. Scalebar= 25μm.

**Supplementary Figure 7 (related to Figure 5). VNC specific *hth[RNAi]* rescues.** Restriction of *hth* knockdown to the *VT-7068* neurons in VNC in *OTD-flp, tub>stop>Gal80 /UAS-hth-RNA[RNAi]; 7068-Gal4, mir-C11/*Δ*mir* flies, is sufficient to revert significantly the elevated egg-laying observed in *mir-iab-4/8* mutants. Mann-Whitney non parametric test, ** p<0.01, **** p<0.0001. Error bars= SEM. Wildtype flies are *Canton-S* strain, *mir-iab-4/8 −/−* mutants=*mir[*Δ*/C11]* transheterozygotes.

**Supplementary Figure 8 (related to Figure 6). Hth engineering pipeline and patterning in *hth-HD* 3’UTR alleles.** (A) CRISPR/ΦC31 approach to generate the 3’UTR engineering platform in *hth-HD* isoform. Hth pattern in the entire abdominal region of larval VNCs (B) and the dorsal region of adult AbG of *hth-HD* 3’ UTR alleles. Scale bar= 50μm.

**Supplementary Figure 9 (related to Figure 7). Virgin indexes of hth 3’UTR engineered alleles.** Both mutation of miR-iab-4/8 binding sites and deletion of the neuronal extension shift virgin behavior to mated PMR.

**Supplementary items:**

**Supplementary movie 1 (related to Figure 1).** Mating assay. Wildtype virgins and wildtype males.

**Supplementary movie 2 (related to Figure 1).** Mating assay. *mir[*Δ*/C11]* virgins and wildtype males

