## Supplementary Figures for "miRNAs and neural alternative polyadenylation specify the virgin behavioral state"

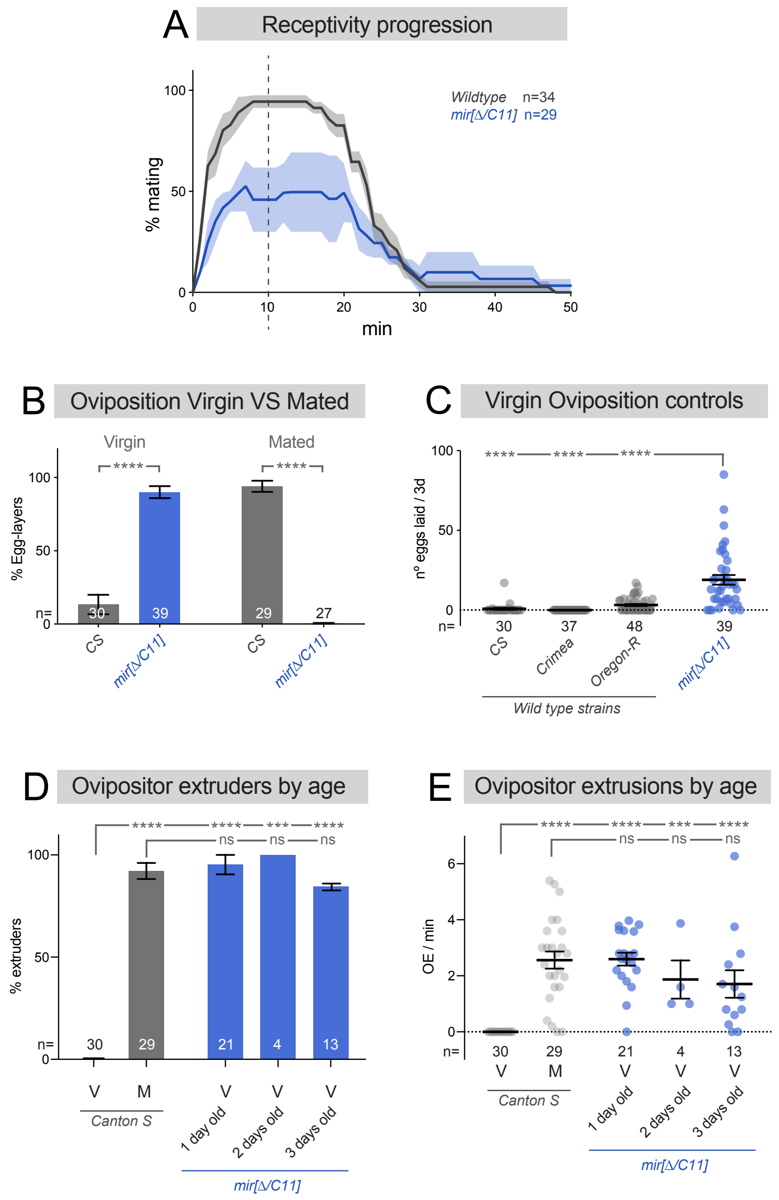


**Supplementary Figure 1 (related to Figure 1). Extended behavioral analysis of *mir-iab-4/8* mutants.** (A) Temporal analysis of sexual receptivity, assessed by the fraction of individual females that are copulating at a given timepoint. The 10 minute timepoint (dotted line) was used subsequently to summarize and compare multiple genotypes or manipulations, shown in Figure 1C. (B) Qualitative oviposition in wildtype and *mir[∆/C11]* flies. Egg-laying is uncoupled from internal state. (C) Quantitative egg-laying counts across several wild type strains. Qualitative (D) and quantitative (E) representation of ovipositor extrusions. Overall levels of male rejection by extruding the ovipositor is maximal in young virgins (1 day old). Fisher's exact test for qualitative analysis (B, D), and Mann-Whitney non parametric test for quantitative analysis (C, E). ns=not significant, * p<0.05, ** p<0.01, *** p<0.001, **** p<0.0001. Error bars = SEM.

Garaulet et al,
Supplementary Figure 1

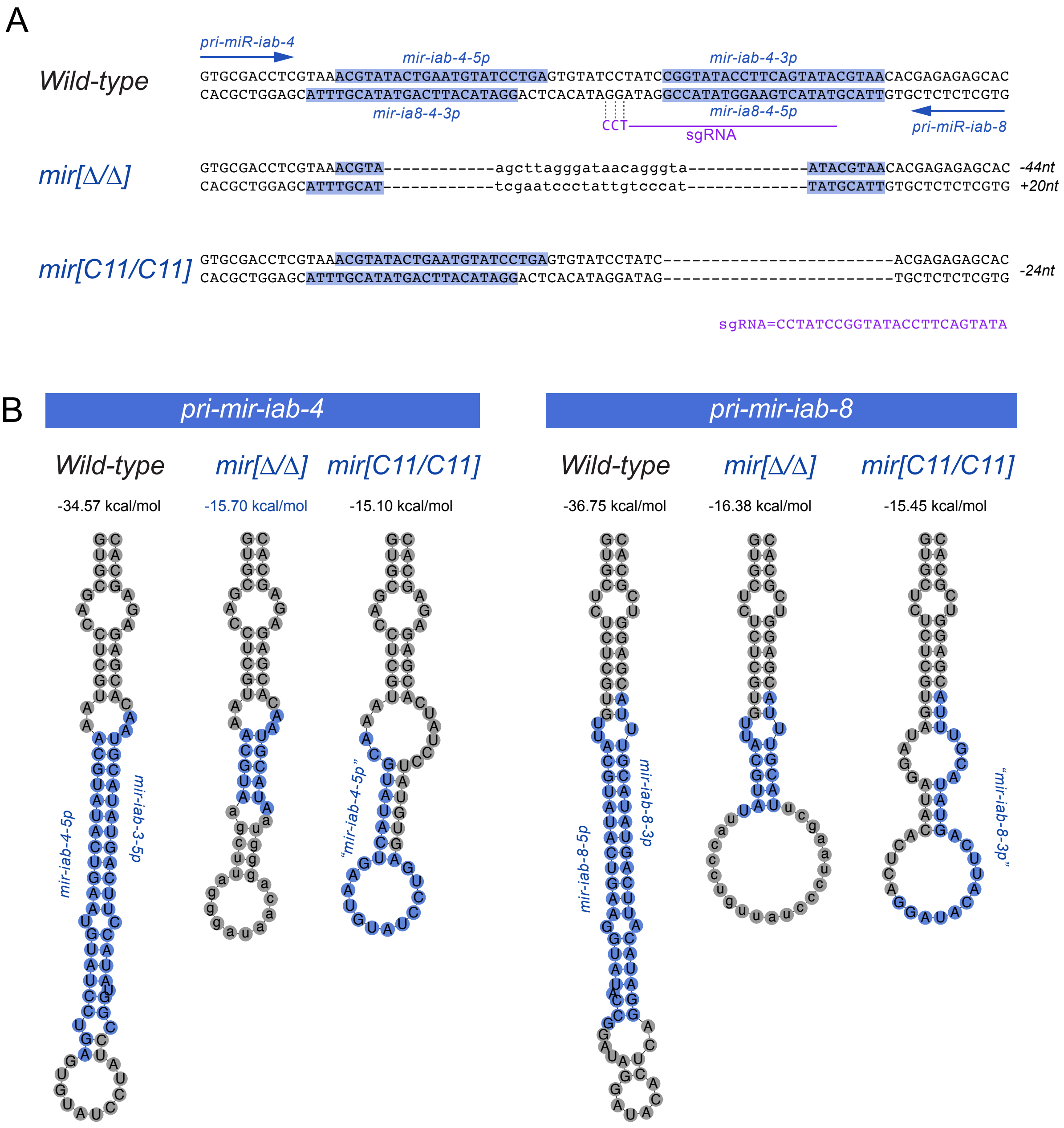


**Supplementary Figure 2. *mir-iab-4/8* mutant alleles.** (A) Wildtype, *mir[Δ/Δ]* and *mir[C11/C11]* sequence. (B) RNAfold predictions of each allele. The newly made *mir[C11]* allele deletes miR-iab-8-5p (miR-iab-4-3p) and does not adopt a pre-miRNA-like hairpin structure. Blue= *mir-iab-4/8* species in each allele. Magenta= sgRNA used to mutagenize *mir-iab-4/8 locus*. Low-case letters: inserted nucleotides. nt=nucleotides.

Garaulet et al,
Supplementary Figure 2

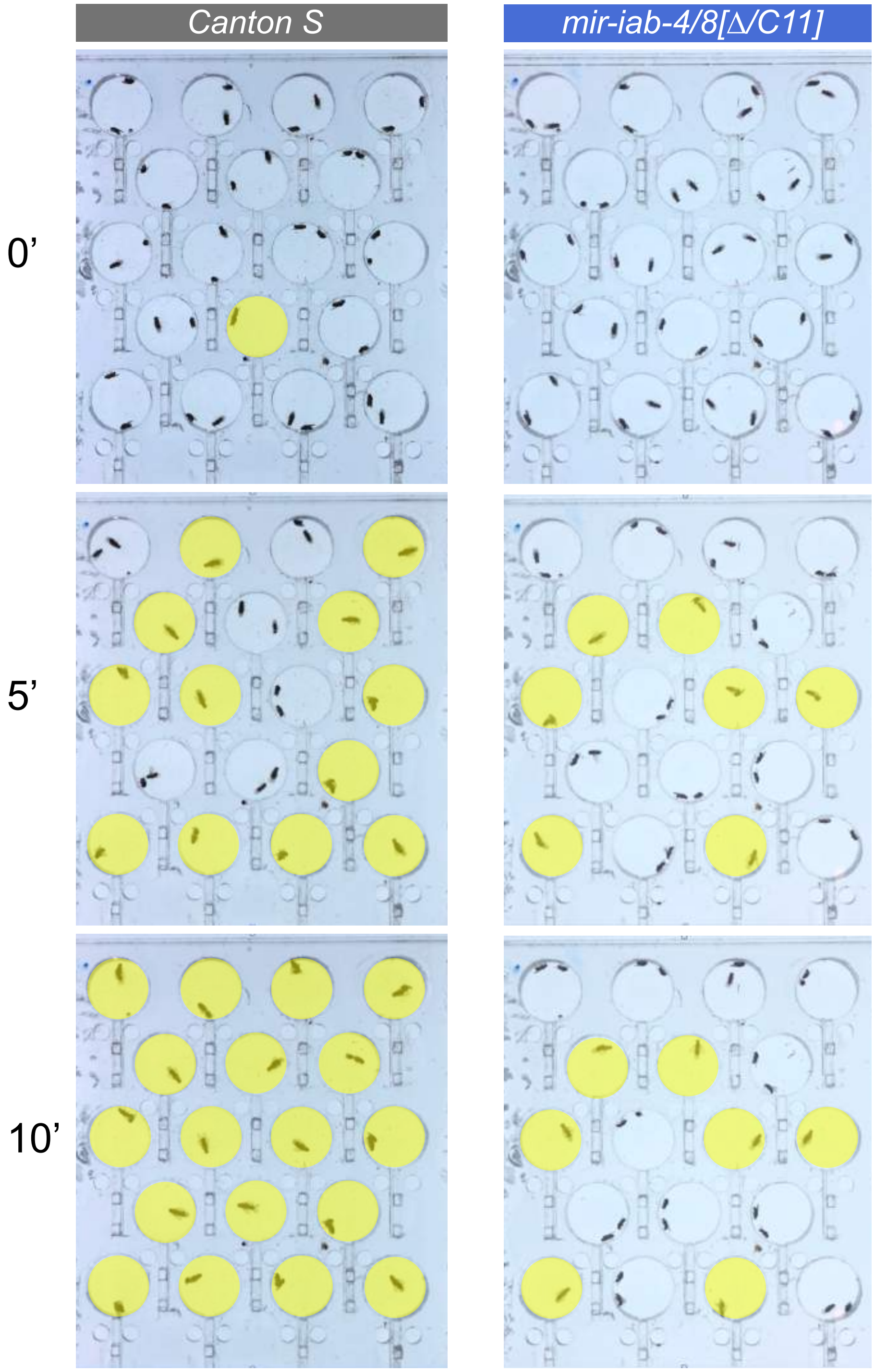


**Supplementary Figure 3. Receptivity dynamics in *mir-iab-4/8* mutants.** Time lapse shots from movies of 18 *wildtype* (*Canton-S*) and 18 *mir[Δ/C11*] female flies. In all cases males are *Canton-S*. Mating couples are marked in yellow. Compared to control, the copulation success of *mir[Δ/C11]* mutants is markedly reduced.

Garaulet et al,
Supplementary Figure 3

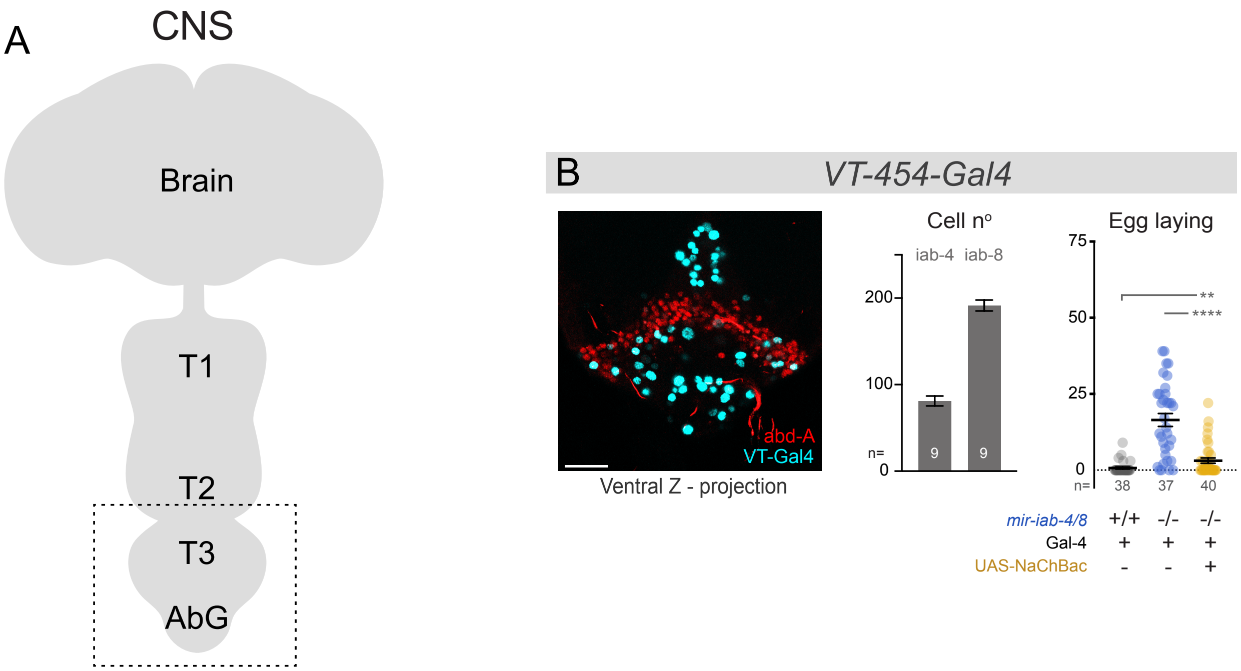


**Supplementary Figure 4 (related to Figure 2). Quantification of additional VT-switch neurons.** (A) Schematic of the Drosophila CNS. Boxed region corresponds to the ventral side of AbG shown in B-E. (B) Z-projection of *454-Gal4* VT-switch neurons (cyan) co-stained with Abd-A (red). Number of VT-switch+ cells per VNC domain (*iab-4* vs. *iab-8*). Egg-laying counts in *VT-454>UAS-NaChBac* virgins. Mann-Whitney non parametric test, ** p<0.01, **** p<0.0001. Error bars = SEM. Scalebar = 25µm.

Garaulet et al,
Supplementary Figure 4

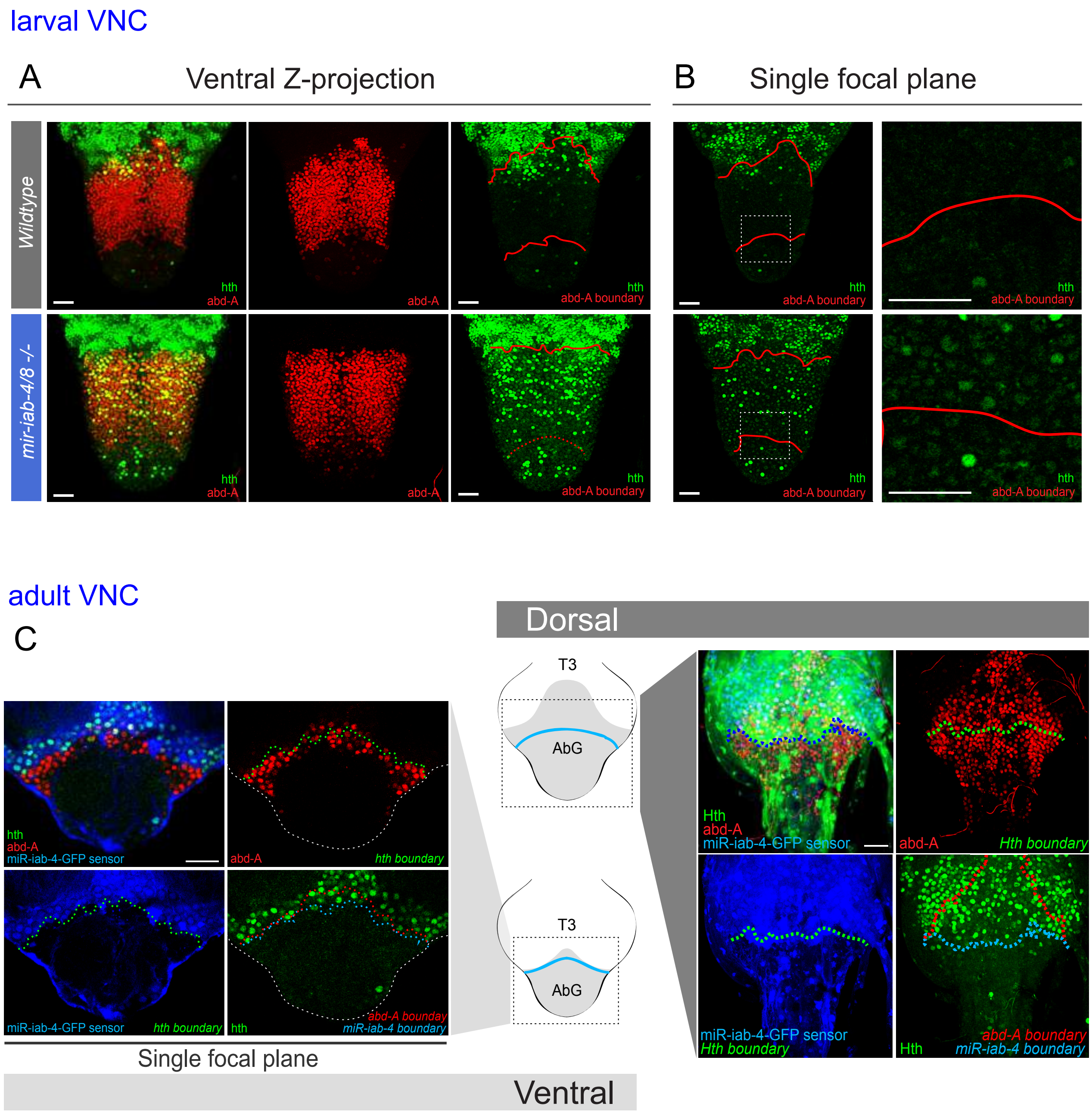


**Supplementary Figure 5 (related to Figure 4). Extended characterization of Hth expression: I.** (A, B) Hth immunostaining (green) in larval VNCs. (A) Hth is derepressed in abdominal segments (Abd-A, red) of *mir[Δ/C11]* female larvae (Z-projections). (B) Mild levels of Hth accumulation are detected in nearly all cells of mutant VNCs. Along the dorsoventral axis, Hth is mutually exclusive with miR-iab-4 and miR-iab-8 (inferred from activity *tub-GFP-sensors*), but not with Abd-A (C). Scalebar= 25μm.

Garaulet et al,
Supplementary Figure 5

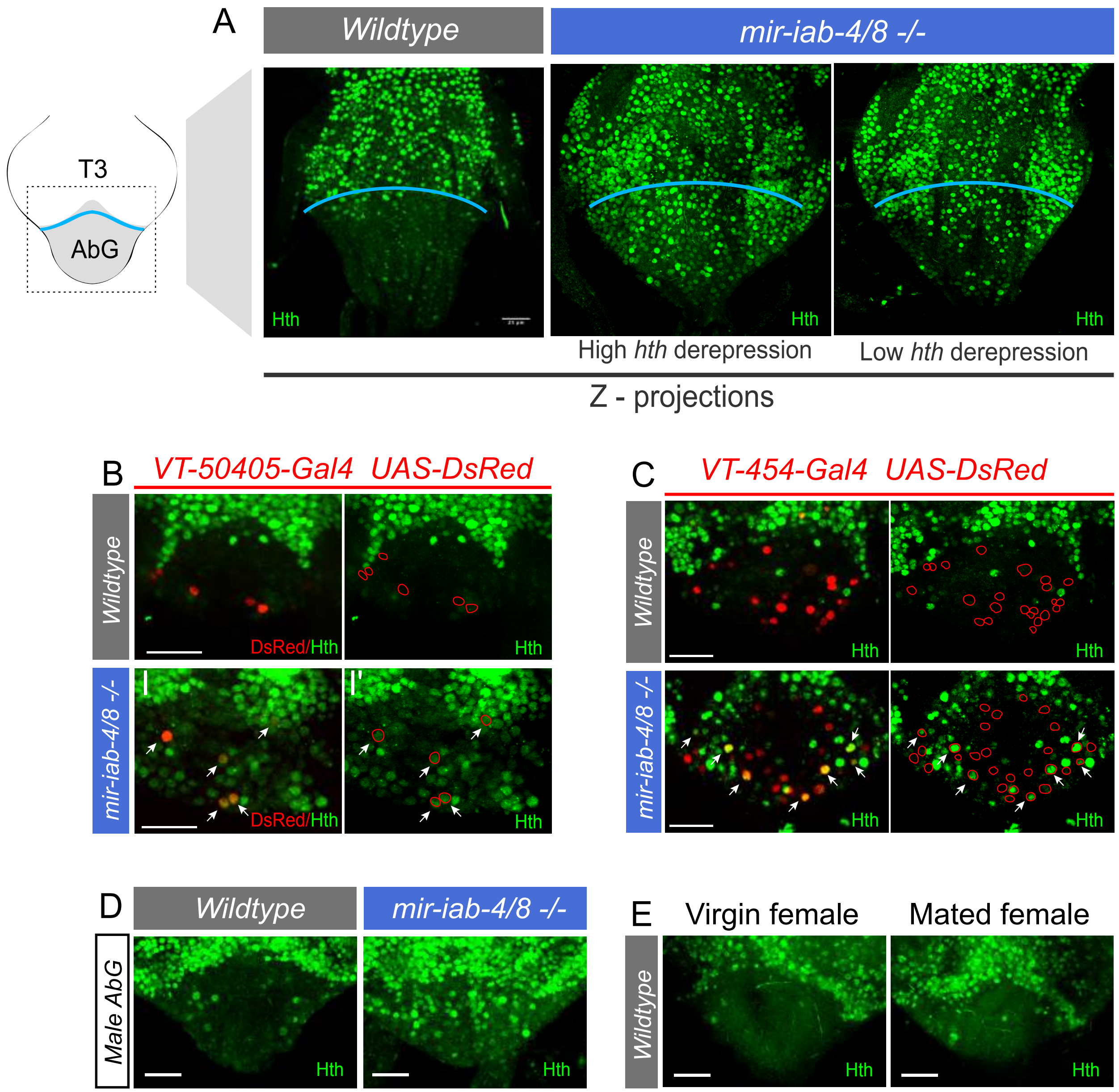


**Supplementary Figure 6 (related to Figure 4). Extended characterization of Hth expression: II.** (A) Hth accumulation in wildtype and two *mir-iab-4/8* mutants, illustrating variability of derepression in mutant VNCs. (B,C) Hth expression relative to VT-switch neuron populations. Only showing the AbG boxed in A. Some *VT-454* labeled neurons show especially high accumulation of Hth protein (arrows). (D) Hth expression in AbG of wildtype and *mir-iab-4/8* mutant males is similar to their female counterparts. (E) Copulation does not affect Hth expression levels in the female AbG. Scalebar= 25μm.

Garaulet et al,
Supplementary Figure 6

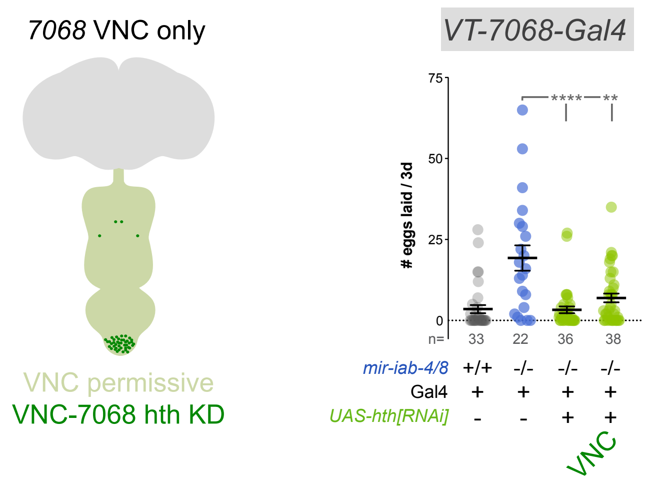


**Supplementary Figure 7 (related to Figure 5). VNC specific *hth[RNAi]* rescues.** Restriction of *hth* knockdown to the *VT-7068* neurons in VNC in *OTD-flp, tub>stop>Gal80 /UAS-hth-[RNAi]; 7068-Gal4, mir-C11/Δmir* flies, is sufficient to revert significantly the elevated egg-laying observed in *mir-iab-4/8* mutants. Mann-Whitney non parametric test, ** p<0.01, **** p<0.0001. Error bars= SEM. Wildtype flies are *Canton-S* strain, *mir-iab-4/8 -/-* mutants=*mir[Δ/C11]* transheterozygotes.

Garaulet et al,
Supplementary Figure 7

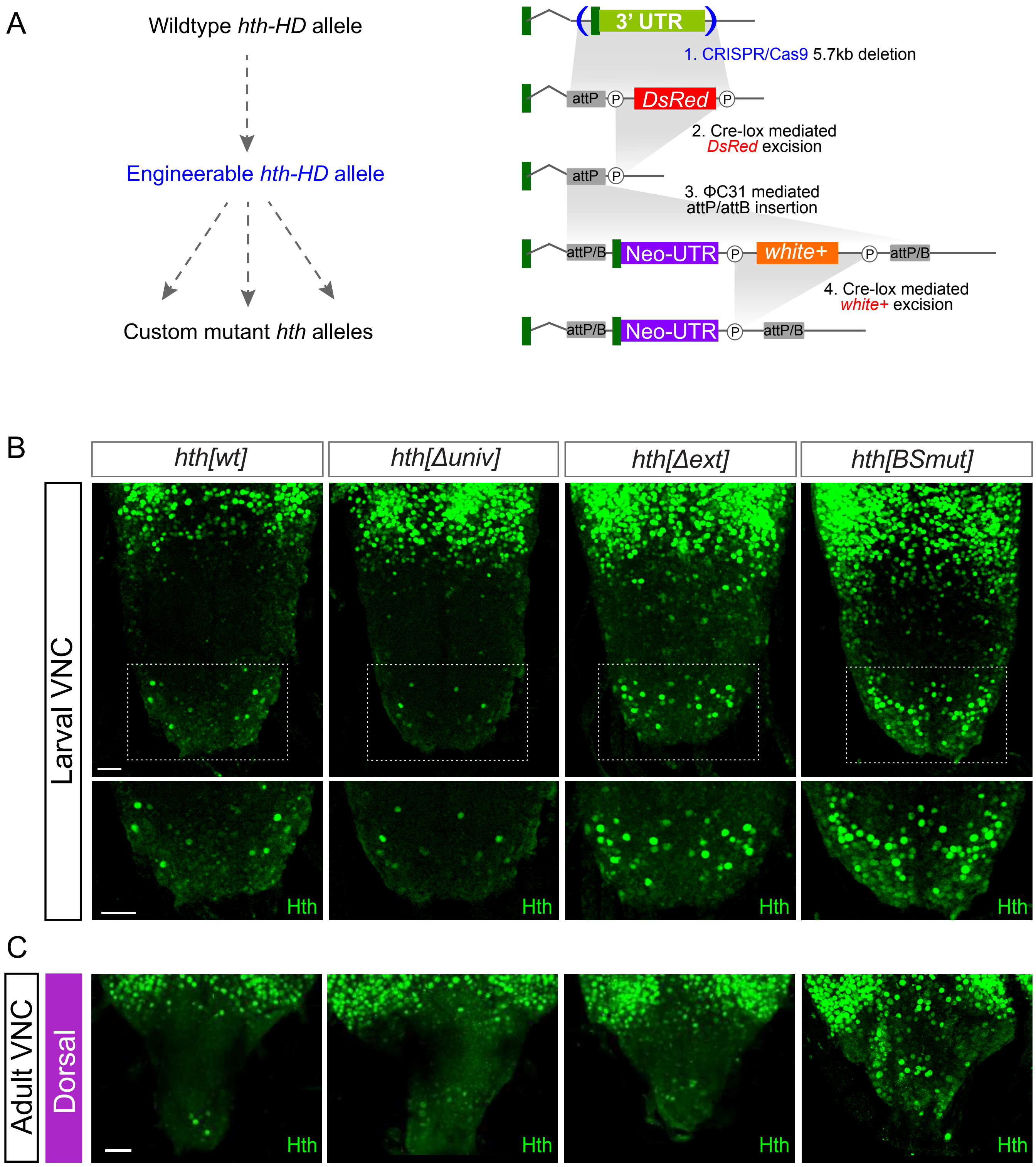


**Supplementary Figure 8 (related to Figure 6). Hth engineering pipeline and patterning in *hth-HD* 3´UTR alleles.** (A) CRISPR/ΦC31 approach to generate the 3´UTR engineering platform in *hth-HD* isoform. Hth pattern in the entire abdominal region of larval VNCs (B) and the dorsal region of adult AbG of *hth-HD* 3' UTR alleles. Scale bar= 50µm.-

Garaulet et al,
Supplementary Figure 8

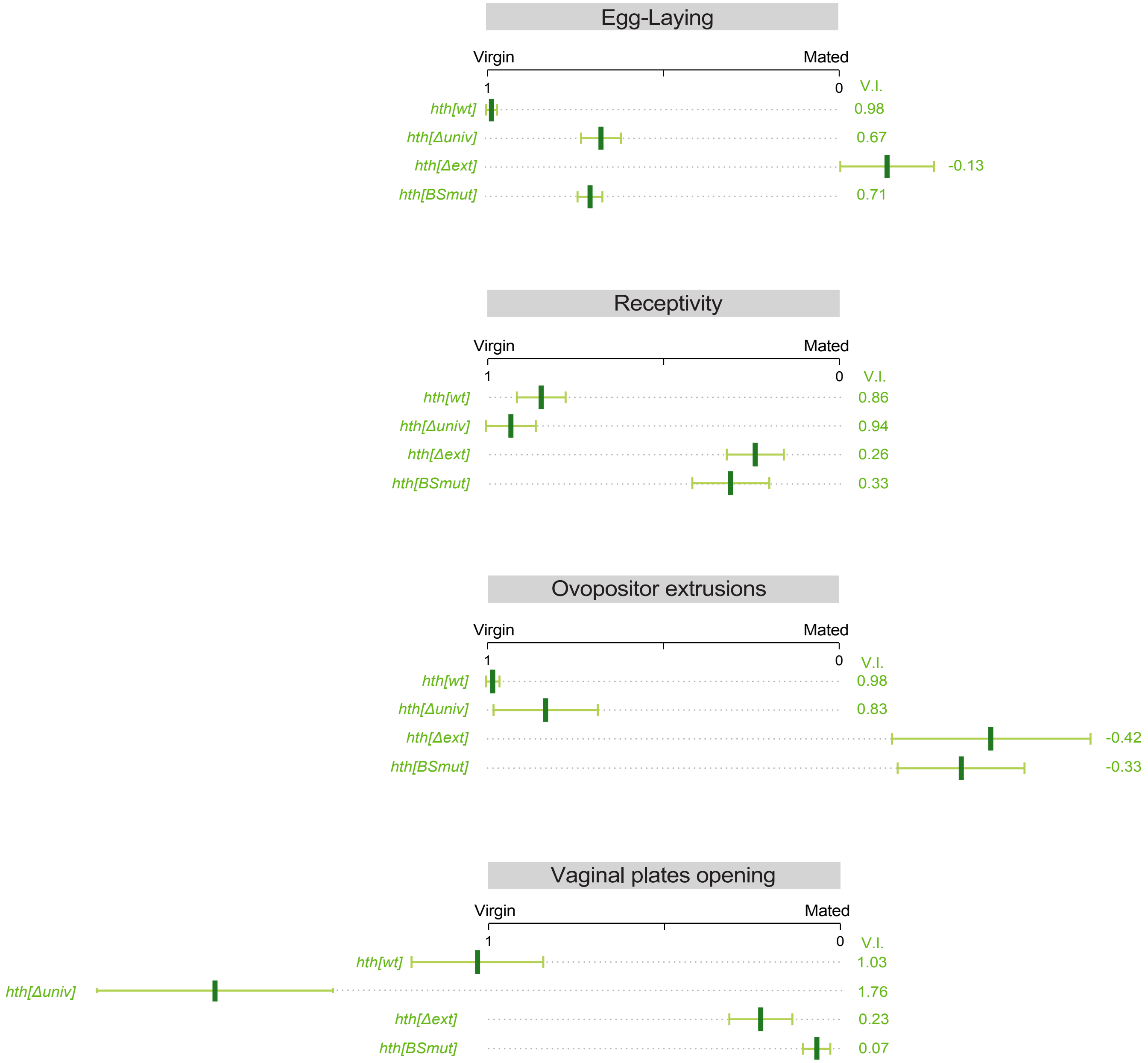


**Supplementary Figure 9 (related to Figure 7). Virgin indexes of *hth-HD* 3'UTR engineered alleles.** Both mutation of miR-iab-4/8 binding sites or deletion of the neuronal extension shift virgin behavior to mated PMR in virgins.

Garaulet et al,
Supplementary Figure 9
